# Unsilenced inhibitory cortical ensemble gates remote memory retrieval

**DOI:** 10.1101/2024.07.01.601454

**Authors:** Shaoli Wang, Tao Sheng, Feng Su, He Yang, Rui Cao, Qiao Wang, Wei-Qun Fang, Chen Zhang, Wei Lu

## Abstract

Acquired information can be consolidated to remote memory for storage but persists in a dormant state until its retrieval. However, it remains unknown how dormant memory is reactivated. Using a combination of simultaneous two-photon calcium imaging and holographic optogenetics in the anterior cingulate cortex (ACC) in vivo, we discover a subset of GABAergic neurons that are specifically associated with dormant memory retrieval. These interneurons display persistent activity and inter-neuronal synchronization at the remote memory stage. In the absence of natural contextual cues, directly activating these interneurons reliably recalls cortical ensembles relevant to remote memory retrieval with context specificity. Conversely, targeted volumetric inactivation of these interneurons suppresses context-induced memory retrieval. Our results reveal an unexpected role of unsilenced inhibitory cortical ensembles in causally gating the retrievability of dormant remote memory.

## Main text

Memories are thought to be initially encoded in the hippocampus (recent memory) but subsequently consolidated within the neocortex for permanent storage (remote memory)^1–6^. Accumulating evidence demonstrates that system consolidation transforms the labile and temporary memory trace to a mature and stable state in the cortex^7–11^. These memory traces undergo constant reorganization during system consolidation^12,13^ and may persist in a dormant state for years until being reactivated by encoding cues^14–18^.

To retain their functions in memory retrieval, memory traces should possess “retrieval handles” that permit activation by specific representations^7,10,19,20^. Memory consolidation may enhance subsequent retrievability of the memory trace, consistent with the idea that “retrieval handles” are established after memory formation^21,22^. To date, however, it remains unknown how dormant traces are awakened for memory retrieval and whether such “retrieval handles” truly exist.

Here, to address this question, we focused on mesoscopic neuronal circuit activities in the anterior cingulate cortex (ACC). This region was found to be critically involved in remote memory storage and receives massive projections from other areas, such as the anterior thalamus, entorhinal cortex, and hippocampus^23–30^. Optogenetic manipulations revealed that these multi-region ACC afferents participate in the successful retrieval of multimodality associative memories^31,32^. The ACC can thus act as an information integration and executive hub for coordinating memory traces. Via longitudinal tracking of neuronal calcium activities, we uncovered persistent activities and inter-neuronal synchronization among a small subset of GABAergic neurons. Via holographic single cellular optogenetic manipulations, we further demonstrated the causal role of inhibitory ensembles in gating remote memory retrieval.

## Results

### Unsilenced ACC neurons at the remote memory stage

We adopted a previously well-characterized behavioral paradigm that enables simultaneous longitudinal two-photon imaging in head-fixed mice (Fig.1a,b)^33,34^. This design aims to temporally align behavioral responses with neuronal activation changes. Briefly, it is a modified contextual conditioning paradigm tailored for head-restricted mice, using a virtual context as the conditioned cue, nociceptive stimuli at the back paw as the unconditioned cue, and detecting "fidgeting" responses with instant pressure changes of the forepaws (see Methods for details). By properly adjusting parameters for pressure detection and mouse training, we ensured that the virtual contextual conditioning reliably induced immediate fidgeting responses (Extended Data Fig. 1g), specifically corresponding to animal’s immobility as inferred by simultaneous video analysis (Fig. 1c). With this setup, we adjusted parameters to induce and maintain remote memory formation and confirmed that mice exhibited obvious fidgeting behaviors when re-exposed to the conditioning context on the retrieval day (day 63; Fig. 1d)^35^. This indicated that the associative memory was successfully maintained and retrieved. It is important to note that the mice were retrieved only once on day 63 post-conditioning to avoid further activation of memory information. As controls, mice trained with conditioning stimuli only (CS-only) or those that were home-caged at all times did not display fidgeting responses to the contextual stimuli.

**Fig. 1.**
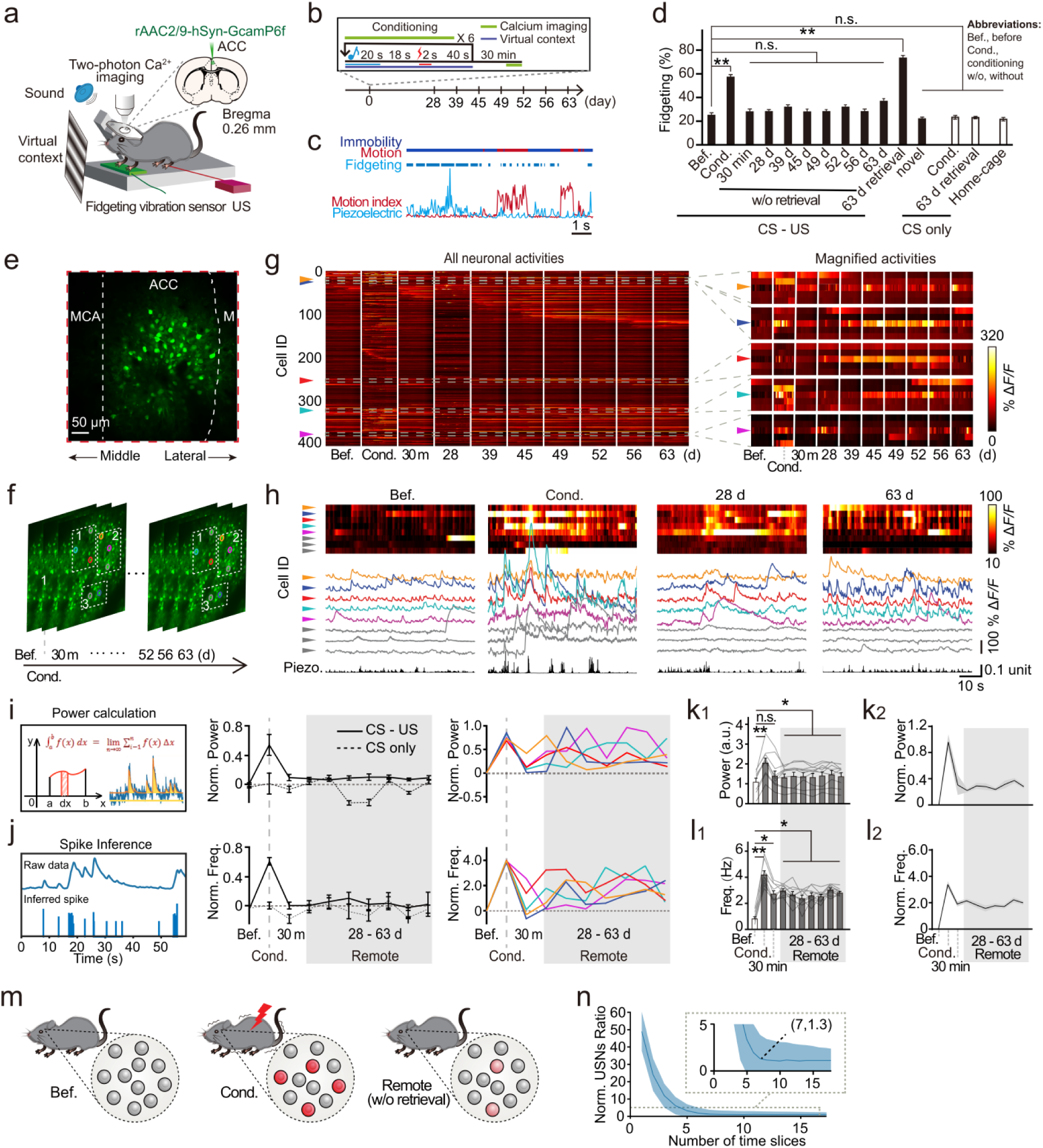
A small number of USNs are identified at remote memory stages. **a**, Schematic diagram of the in-vivo imaging setups. US, unconditioned stimuli. **b**, Experimental timeline. **c**, Validation of conditioned behavioral response assessed by video analysis and piezoelectric signals respectively. Dark blue, period of immobility; Red, period of motion. Raw piezoelectric signals (light blue) and raw video analyzed motion index (red) were in reverse of each other since they respectively correspond to amounts of immobility and motion. Piezoelectric sensors pick up immobility since animals’ paw pressure intensifies during the freezing response. Data was from the last training trial. **d**, Successful formation and retrieval of context (conditioned stimuli, CS) foot shock (unconditioned stimuli, US) associative memory. The CS-US group showed a significantly higher fidgeting response during remote memory retrieval (day 63) compared to all other time points (without retrieval), *P* < 0.0001. CS-only and home-caged controls showed no changes, *P* > 0.1, one-way ANOVA with Tukey’s post hoc test, \**P* < 0.05, \*\**P* < 0.01, mean ± s.e.m. Conditioning data were from the last conditioning trial as in all the following figures. “Bef.” in the plot refers to before conditioning. **e**, Imaging window position at the ACC (AP: 0.26 mm, ML: -0.35 mm, DV: 0.30 mm). Middle cortical artery (MCA) was used as a reference point. **f**, Schematic diagram of two-photon images at a depth of 200-300 μm. Colored circles mark 5 neurons with persistently increased activities above the before-conditioning level. Each color denotes cellular ID, consistent in all following figures g-j. Gray circles mark 3 randomly selected neighboring neurons with activities fluctuating around the baseline. **g**, Heatmaps showing calcium transients (ΔF/F) from all identified neurons within the imaging field at different imaging time points. Right, magnified images in dotted lined boxes on the left. Among all the detected neurons, 5 of them (ID marked by arrow colors) exhibited persistently increased activities at the remote memory stage (post-conditioning day 28-63). The color bar indicates ΔF/F values. n = 405 neurons from 1 animal. **h**, Temporally aligned activity heatmap (top), raw calcium traces of selective neurons (middle), and fidgeting response (bottom) obtained at different time points. **i**, Sustained increase in USNs during the entire remote memory stage revealed by calcium power analysis. The left diagram shows the power calculation method. The line charts at right show temporal changes in normalized neuronal calcium power in all neurons (black line), USNs (colored line), or randomly selected neighboring neurons (grey line). The dotted line at zero indicates the normalized before-conditioning baseline. The shaded portions indicate the remote memory stage. Normalization was performed by subtracting the average calcium power of the before-training baseline from the average power of each time point. **j**, Neuronal spike counts based on spiking activity inference algorithms identified the same USNs. The diagram shows reconstructed neuronal calcium spikes obtained by the deep learning method DeepSpike. Calibration bar: 5 s, 40% ΔF/F. The arrangement of line charts matched those in panel I. **k,l**, Bar graphs showing modestly elevated calcium power (k) and spike frequencies (l) in USNs during the remote memory stages (k1, Bef. vs. Cond. P < 0.001; Bef. vs. Remote, P < 0.05; l1, Bef. vs. Cond. *P* < 0.01, Bef. vs. 30 min, *P* < 0.05, Bef. vs. Remote, *P* < 0.01, each timepoint’s value was compared to Bef.’s value using nonparametric Friedman test with Dunn’s correction for multiple comparisons, n = 41 neurons from 11 animals). k1 and l1 show non-normalized values; k2 and l2 show values normalized by subtracting data from the before imaging timepoint. These USNs displayed similar silencing of neuronal activity after conditioning. However, their activities did not return to before-conditioning baseline level. Instead, they maintained their activities at a modest level above baseline throughout the remote stage. Power a.u., power in arbitrary unit (a.u.). Gray area covers remote time points. **m**, The diagram illustrating the neuronal activity evolvement across different stages. Note that at the remote memory stage, spontaneous activity of USNs does not return to baseline level. Instead, it maintains in an intermediately potentiated state. The weight of red color denotes the neuronal activity level. **n**, Influence of the number of imaging time points on USN identification. When the number of imaging time points (time slices) exceeded 7, the normalized number of active neurons became stabilized (4.33 ± 0.22, n = 11 animals; ratio over total neuronal number shown in the magnification box), justifying our inclusion of 7 remote-stage imaging time points to delineate USNs.

We then tracked neuronal activity dynamics via longitudinal two-photon calcium imaging. We expressed a genetically encoded Ca^2+^ indicator (GCaMP6f, using the viral tool rAAV2/9-hsyn-GCaMP6f) in the ACC (Extended Data Fig. 2a,b, AP: 0.26 mm, ML: -0.35 mm, DV: 0.30 mm) and captured GCaMP6f signals of neurons at a depth of 200-300 μm at 30 Hz through a cranial glass window above the ACC of awake, head-fixed mice continuously for 9 weeks (Fig. 1a,e,f)^36,37^. Employing the middle artery as a reference, we precisely located the ACC in the imaging planes^38^ to avoid accidentally imaging the neighboring motor cortex (Fig. 1e, Extended Data Fig. 3b,c). The imaging window size was set at 565×565 μm². For each imaging timepoint we selected one of four 90s segments imaged as the final data. The overall neuronal activity level (as indicated by the somatic fluorescence change ΔF/F) increased during the conditioning phase, quickly decreased to the pre-conditioning baseline level after conditioning, and maintained a low activity level during the remote memory phase (28-63 days post-conditioning) (Fig. 1g). However, we unexpectedly observed a small number of neurons (2-6 cells) exhibiting fluctuating but persistently higher activity levels than the baseline during the entire remote memory phase (Fig. 1g,h).

These incompletely silenced neurons drew our attention, as neurons activated during encoding have been proposed to enter a dormant state at the remote memory stage^5,26^. To further clarify their activity patterns, we inferred neuronal spiking events from their ΔF/F transients using two independent quantitative approaches: calcium power calculation (Fig. 1i,k)^39^ and spike inference analysis (Fig. 1j,l)^40^ (see Methods for details). Both assessments yielded consistent results across different subjects, revealing a small subset of neurons with unique activity dynamics over time. Initially, these neurons showed similar silencing of activity approximately 30 minutes after conditioning. However, during the entire remote memory stage (shaded regions in Fig.1i-l), their spontaneous activities did not completely return to pre-conditioning baseline levels. Instead, their activities were maintained at an intermediate level, which was above baseline levels but still obviously lower than that during conditioning stage (Fig. 1k,l). Therefore, their spontaneous activities were insufficient to elicit memory retrieval until further increased upon retrieval at day 63 (Fig. 1k-m; Extended Data Fig.4). We thus defined “unsilenced neurons (USNs)” as the neurons that maintained above-zero normalized calcium power at every imaging time point throughout the remote memory stages (28-63 days; also see Methods for details on quantitative definition).

To avoid false identification of these persistently active neurons due to inadequate sampling times, we calculated their percentage using different numbers of imaging time points. Our data show that when over 7 imaging time points were employed, the percentage of neurons with above-zero activity at the remote memory stage became relatively invariant (Fig. 1n). Comparing neuronal activities at the remote stage in the CS-US and CS-only groups demonstrated that USNs could only be identified in trained animals (Extended Data Fig.5a-e). This finding was further corroborated by bootstrapping neuronal activities across time (Extended Data Fig. 5f,g).

Long-term overexpression of GCaMP sensors in neurons often leads to higher fluorescent signals that don’t necessarily reflect increased neuronal activity, especially in unhealthy neurons. These USNs exhibited suppression of activity after conditioning, similar to most other neurons, and could be successfully photo-activated by two-photon optogenetics at 63 days post-conditioning, indicating their overall health despite long-term GCaMP6f expression (Extended Data Fig. 1c-f). Furthermore, we did not detect any USNs in the neighboring cortical region (e.g., motor cortex, Extended Data Fig. 3d), suggesting that the USNs were indeed located in the surface ACC region and potentially play important roles in remote memory maintenance.

### Unsilenced ACC neurons are GABAergic

We further investigated whether the unsilenced neurons were excitatory or inhibitory. To this end, we crossed Viaat-cre (Jackson Laboratory, strain No: 017535) mice with Ai9 mice (Jackson Laboratory, strain No: 007909) to obtain GABA+/Cre;Ai9wt/mut (GABA-Cre::Ai9) mice, which express mCherry red fluorescent proteins in GABAergic neurons (Fig. 2a)^41^. All the mCherry-expressing neurons were also positive for the interneuron marker glutamic acid decarboxylase 67 (GAD67; Extended Data Fig. 6)^42^, indicating they were indeed GABAergic interneurons. This allowed us to distinguish excitatory neurons from inhibitory neurons with mCherry expression.

**Fig. 2.**
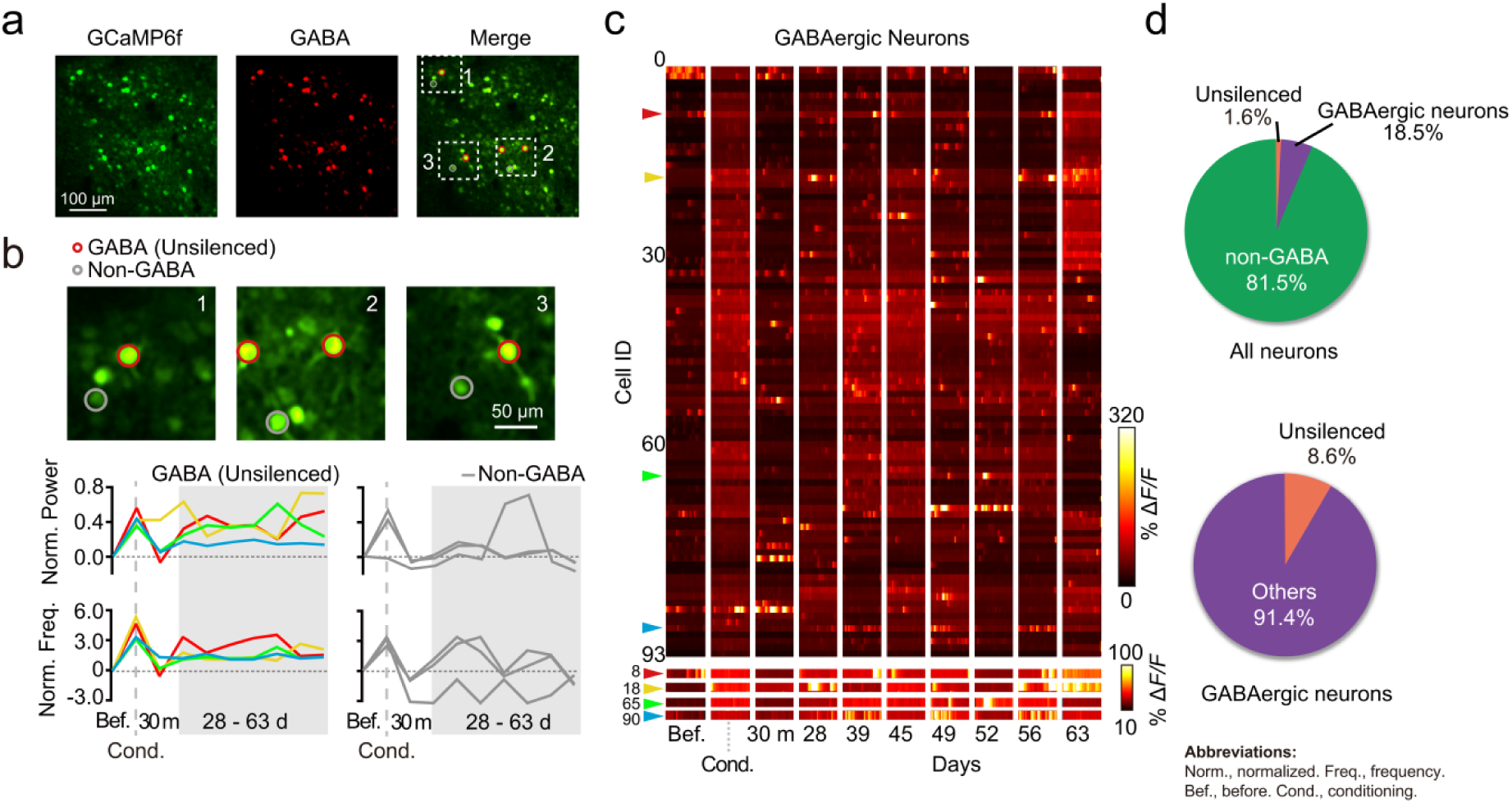
USNs are GABAergic. **a**, Sample image (all time points averaged) in GABA-Cre::Ai9 mice. The left image refers to all GCaMP6f-labeled cells in the imaging field. GABAergic neurons were those labeled by mCherry signals (middle). In merged image (right) neurons in yellow refer to co-labeled GABAergic neurons. Dotted line boxes illustrate the location of USNs (circled in red). **b**, Magnified images marked by dotted line boxes in Fig. 2a demonstrate that USNs are GABAergic. The USNs (circled in red; top), which maintained increased activity (middle, normalized calcium power; bottom, DeepSpike inferred spike frequencies) at the remote stage (post-conditioning day 28-63), were all GABAergic neurons. Neighboring non-GABAergic neurons (circled in gray, activity in gray lines) were employed as the control. **c**, Heatmaps of neuronal activities of all GABAergic neurons recorded across different time points. The USNs maintain increased activity during the entire remote stage. Arrow colors marked cell ID corresponding to (b). Color bar shows ΔF/F value. **d**, Pie chart showing a low percentage of USNs in all imaged neurons and in all GABAergic neurons. The percentage of USNs was calculated by dividing the count of USNs by the total number of all GCaMP+ neurons (n = 11 animals).

We followed the same procedure and criteria to label and identify the USNs in the ACC as described above. All the USNs (Fig. 2a,b) were also mCherry positive, indicating they were GABAergic interneurons. These inhibitory USNs all demonstrated above-baseline activities during the remote stage, while non-GABA controls showed varied activities (Fig. 2c). Among all neurons identified by GCaMP6f expression in the imaging window, we identified 18.5% as GABAergic across all animals. Notably, all identified USNs fell into the GABAergic category, which is 8.6% of the whole GABAergic population and 1.6% of the total population (Fig. 2d). Our findings indicated the existence of a small group of GABAergic USNs, whose activities were induced by the conditioning cues and persistently maintained at the moderately high level during the remote memory stages. This raises important questions: Are these neurons functionally connected? Are they crucial or required for maintaining remote memory?

### USNs maintain highly synchronized activity

To address the above questions, we examined whether the unsilenced GABAergic neurons were functionally connected, as neurons with highly correlated activity can form memory-encoding neuronal ensembles. We based our analysis on the pairwise correlation of spontaneous neuronal activity^27,43^, which were estimated by the time-course calcium spikes inferred from raw calcium ΔF/F traces^44^. We generated weighted adjacency matrices for all mCherry-positive neurons over time (Fig. 3a; see Methods for details). These matrices represented the development of neuronal functional connections over time and were further illustrated as graphs of coactivity complex networks (Fig. 3b). It was clear that the interconnectivity of USNs (red nodes) was initially weak before conditioning, strengthened by conditioning, and then quickly decreased (Fig. 3b). These sequential dynamic connectivity levels were similar to other neurons (gray nodes) and the overall connection of all GABAergic neurons (Fig. 3c). However, during the remote memory stage, the functional interconnectivity of USNs was strengthened and maintained (Fig. 3d,e) across all imaged subjects (n = 6 animals) (Fig. 3g). Therefore, USNs showed enduring high inter-neuronal correlation of spontaneous activity. This indicates that they form a functional ensemble distinct from the entire GABAergic neuron population (Extended Data Fig. 7c).

**Fig. 3.**
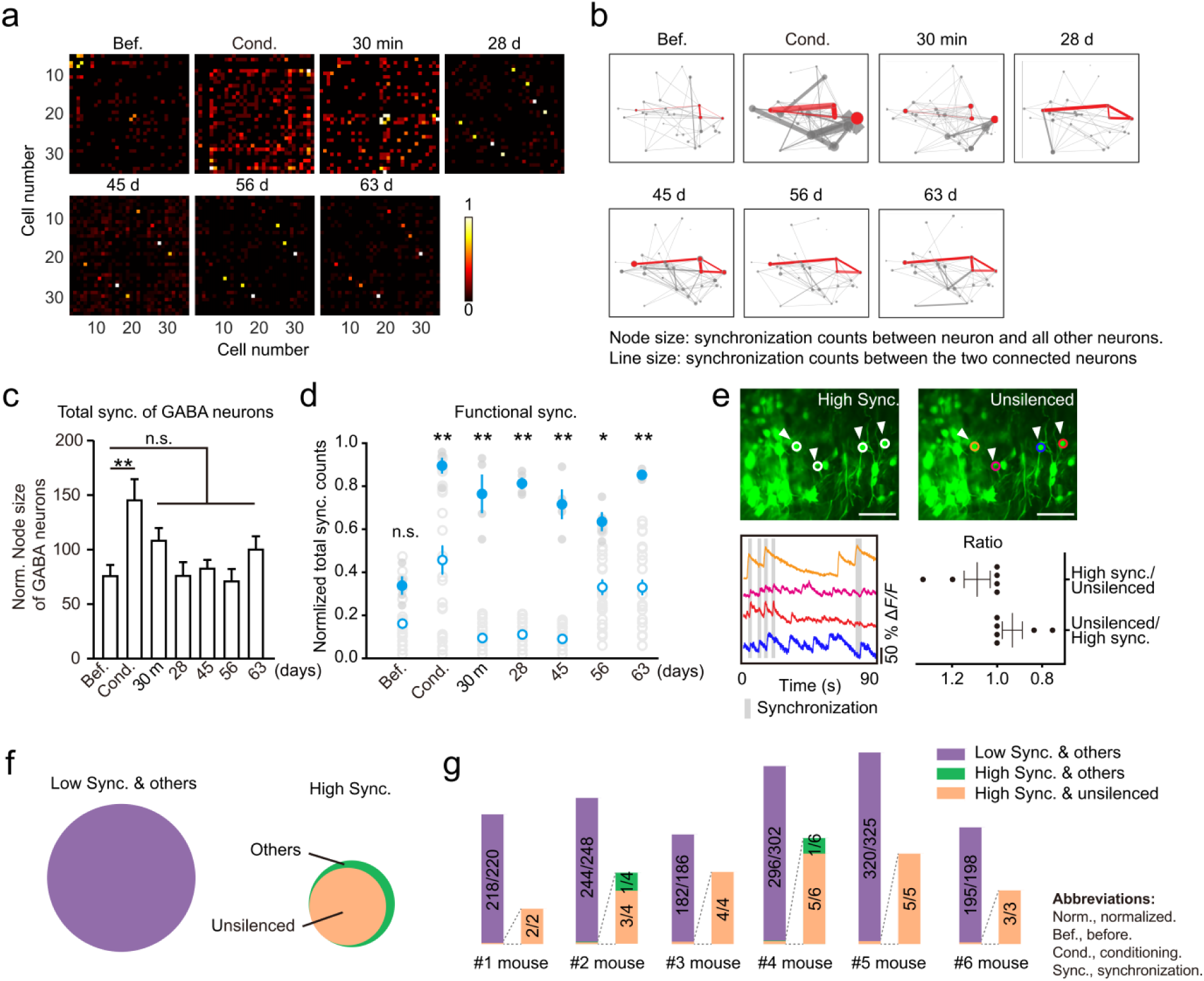
USNs maintain a high synchronization level. **a**, Pairwise synchronization counts between all GABAergic neurons mirrored along the diagonal line. Values were calculated from the temporal co-occurrence of neuronal spikes inferred by the deconvolution algorithm (see Methods). An overall increase in pairwise correlations was detected during the last trial of conditioning, followed by a rundown toward the baseline level. In contrast, heightened pairwise synchronization was maintained between a few GABAergic neurons. **b**, Coactivity complex network of neurons in Fig. 3a. Each node position corresponds to the actual neuronal location in the imaging field. Red, USNs; gray, other neurons. The thickness of the line positively correlates to the synchronization counts between the two neurons. The size of the node positively correlates to the sum of pairwise synchronization counts between the neuron and all other neurons. At the remote memory stage (post-conditioning day 28-63), node size of USNs decreased, while their inter-neuronal edge size increased. This indicates that USNs mostly synchronize with each other at the remote memory stage, instead of maintaining nonselective interactions with all other neurons. **c**, Pairwise synchronization within all GABAergic neurons increased during conditioning and decreased toward the before-training level at the remote memory stage. The node size of GABA neurons refers to the total synchronization counts from all GABAergic neurons. Cond. vs Bef.: *P* < 0.01, one-way ANOVA with Tukey’s post hoc test, \**P* < 0.05, \*\**P* < 0.01, n = 6 animals. **d**, Pairwise synchronization counts yielded two distinct neuronal populations with high synchronization (High Sync; counts ≥ 3σ, filled circles) and low synchronization (Low Sync; counts < 3σ, empty circles). \**P* < 0.05, \*\**P* < 0.01, n = 4 neurons for High Sync. neurons, n=31 neurons for Low Sync. neurons in 1 animal. **e**, Highly matched neuron identity between USNs and High Sync. neurons. Top, representative two-photon images of High Sync. neurons (white circle, left) and USNs (colored circle, right). Bottom left, calcium traces of neurons circled in top right images. Trace colors match with the top right image. Black dots mark calcium spikes (threshold: ΔF/F ≥ 2σ) that synchronize with those of other traces. Bottom right, count ratio of High Sync. neurons and USNs. (unsilenced/High Sync. 0.93 ± 0.05, High Sync./unsilenced 1.09 ± 0.06, n=6). **f**, Venn diagram of USNs and High Sync. neurons. USNs (GABAergic) made up most of the High Sync. neurons. Others refer to cells with calcium power that fluctuate or drop below baseline activities during remote stages. **g**, Distribution of the numbers of unsilenced and High Sync. neurons across different subjects.

Could the unsilenced GABAergic neurons constitute a stable ensemble throughout the remote memory phase? Answering this question is crucial for understanding whether these neurons are involved in remote memory encoding and processing. To address this, we assessed the stability of USNs’ activity patterns during the remote memory stage by extracting their topological structures (Extended Data Fig. 8a,b) and calculating their joint information entropy (Extended Data Fig. 8c; see Methods for details)^45–47^. Compared to randomized neuronal groups, ensembles of USNs exhibited a sustained decrease in average joint information entropy (Extended Data Fig. 8e) and an increase in average joint mutual information (Extended Data Fig. 8f) starting from the beginning of the remote memory stage at P28. Furthermore, decreased entropy was observed only in USNs, not in the total GABAergic neuronal population or the entire imaged population (Extended Data Fig. 9). Additionally, we used various analyses, including comparisons to shuffled surrogates, to ensure that the elevated "synchronization" between USNs (Extended Data Fig. 10) and the reduced joint information entropy were not due to increased firing rates (Extended Data Fig. 8d; see Methods for details). We further calculated Mahalanobis distance between activity matrices of remote stage and pre-conditioning baseline (Extended Data Fig. 8g,h) to confirm that USNs exhibited convergent temporal activity dynamics that were different from other GABAergic neurons^48^. Collectively, these results suggest that USNs constitute stable functional networks, potentially serving as substrates for remote memory maintenance and future retrieval.

### Targeted activation of USNs retrieved remote memory

Our findings that USNs were exclusively GABAergic and formed a stable ensemble are both exciting and surprising, as they seem to contradict the traditional view that memory traces are primarily driven by excitatory neuronal activities^31^. To directly test the possible involvement of these unsilenced GABAergic ensembles in remote memory retrieval, it is necessary to selectively activate them during the remote memory stage and examine the possible following effects on memory retrieval. For this purpose, we performed simultaneous imaging and optogenetics using a holographic two-photon microscope equipped with two lasers: one for imaging GCaMP6f at 920 nm and the other for photo-activation of the redshifted opsin C1V1 at 1,035 nm^49–53^. By co-expressing GCaMP6f and C1V1 in the same neurons, we were able to identify and subsequently photo-activate the USNs (Fig. 4a). The constancy, robustness, and precision of photo-activation was tested by sequential photo-stimulation at the soma of selected C1V1-expressing neurons with variable intervals (see Methods for details), resulting in obvious depolarization of the targeted neurons without evoking activation of neighboring non-targeted neurons (Fig. 4b; Extended Data Fig. 11a,b).

**Fig. 4.**
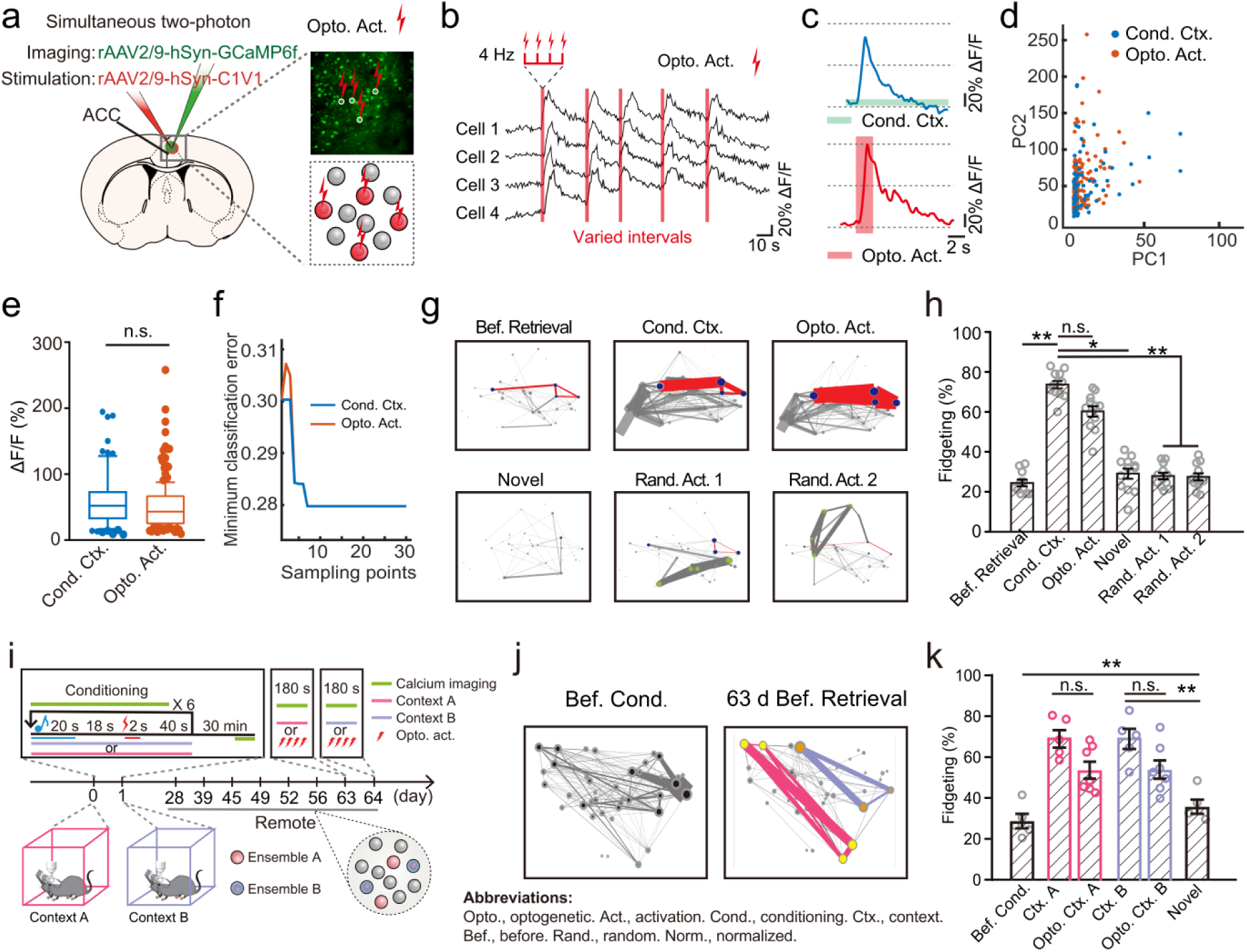
Holographic photo-activation of USNs retrieves remote memories with context specificity. **a**, Experimental schematics. Single-cell resolution stimulation was achieved via either a spatial light modulator (SLM) or rapid movement of the galvo-galvo mirrors. **b**, Validation of holographic stimulation. Holographic stimulation evoked robust and repeatable elevated calcium transients in randomly selected test neurons. The time width of the optogenetic activation (Opto. Act.) pulse is 2 seconds, and the illumination frequency within the pulse is 4 Hz. To better test the stimulation efficacy, the optogenetic stimuli were delivered with unfixed interval. For the retrieval experiment, however, each stimulation was delivered 5 times at 10 s apart. **c**, Comparable calcium response level of an example USN during context retrieval and Opto. Act. **d**, Overlapping principal component distribution of average ΔF/F response of USNs (n=5 animals) under Cond. context (Cond. Ctx; blue) and Opto. Act. (red). **e**, No significant difference between the average ΔF/F of USNs (n = 5 animals; Cond. Ctx = 60.19 ± 4.94 %, Opto. Act. = 51.58 ± 2.79 %, *P* = 0.16, unpaired t-test.) **f**, We trained a decision tree classifier based on results in e, Cond. Ctx and Opto. Act. evoked similar activities among USNs, resulting in a similar distribution of minimal classification error. X-axis denotes the number of samples used in training. **g**, Similar coactivity network patterns upon optogenetic and conditioning contextual memory retrieval. As the control, novel contextual cues or stimulation of randomly selected GABAergic neurons elicited completely different coactivity patterns. Blue nodes, USNs; green nodes, randomly stimulated GABAergic neurons. Red edges denote pairwise synchronization between USNs. For node and wedge calculations, see Fig. 3b. **h**, Successful remote memory retrieval by optogenetic stimulation of USNs. Optogenetic stimulation elicited a similar fidgeting rate to that induced by conditioning contextual stimulation (*P* = 0.125 *>* 0.05; n = 12, one-way ANOVA with Tukey’s post hoc test, mean ± s.e.m.). In contrast, the novel context or activation of the same number of randomly selected neurons elicited a much lower fidgeting rate (random activation: Rand. Act., Rand. Act. 1 *vs.* Bef. Retrieval, *P* = 0.766 > 0.05, Rand. stim2 vs. Bef. Retrieval, *P* = 0.839 > 0.05, Rand. Act. 1 *vs.* Novel, *P >* 0.05, Rand. Act. 2 *vs.* Novel, *P =* 0.999 *>* 0.05; n=12, one-way ANOVA with Tukey’s post hoc test, mean ± s.e.m.). **i**, Experimental setup of the two-context experiment. Mice underwent context A and context B conditioning respectively with 24 hours apart; each memory was then tested 63 d post conditioning respectively. **j**, Coactivity network patterns depict the formation of two context-specific subsets of USNs. Each subset of USNs was determined by checking whether calcium power was persistently above their respective before-training baseline activities for every remote-stage imaging time point. Distinct subsets of USNs (marked by yellow and orange) correspond to the two contexts (pink for context A and purple for context B). **k**, Successful retrieval of remote memories by optogenetic stimulation of USNs corresponding to their respective contexts (Ctx. A *vs*. Bef., *P* < 0.01; Ctx. B *vs*. Bef., *P* < 0.01; Opto. Ctx. A *vs.* Bef., *P* < 0.01; Opto. Ctx. B *vs.* Bef., *P* < 0.01, one-way ANOVA with Tukey’s post hoc test, n = 7 animals, mean ± s.e.m.). Optogenetic activation of each of the two subsets of USNs elicited similar fidgeting responses as their corresponding conditioning contexts, indicating that targeted activation of these cortical neurons is sufficient to retrieve remote memory (Ctx. A *vs.* Novel, *P <* 0.01, Opto. Ctx. A vs. Novel, *P* = 0.006 *<* 0.01; Ctx. B *vs.* Novel, *P <* 0.01, Opto. Ctx. B *vs.* Novel, *P =*0.005 *<* 0.01, one-way ANOVA with Tukey’s post hoc test, n = 7 animals, mean ± s.e.m.).

To further assess the effect of unsilenced neuronal ensemble activation, we performed simultaneous multi-neuron photo-stimulation using a spatial light modulator (SLM) to split a laser beam into multiple beamlets, precisely stimulating selected neurons (Extended Data Fig. 11c). The photo-activation effect was confirmed by comparing the ΔF/F response of USNs under contextual recall and optic stimulation via principal component analysis and decision tree classifiers (Fig. 4c-f). The cellular response under optogenetic stimulation highly resembled the natural cued effects, justifying the robustness of holographic light stimulation (Fig. 4e). At 63 days after conditioning, simultaneous photo-activation of all USNs (Fig. 4h; see Methods for details) lead to obvious fidgeting movements (Fig. 4c), suggesting successful memory retrieval. Consistently, photo-activation rapidly evoked neuronal ensemble activation patterns that were highly similar to those elicited by the conditioning context (Fig. 4d). Previous studies have shown that reactivation of memory-related engram cells can retrieve memory^54^, and optogenetic reactivation was generally conducted on large populations (estimated to be hundreds) of cells^11,31^. To determine if the retrieval of memory-like behavior is simply elicited by artificial photo-activation of a subpopulation of engram cells, we identified memory-related GABAergic cells that were substantially activated during conditioning and conducted optogenetic stimulation on random subsets of these cells (see Methods for details). We found that random reactivation of the same number (2-6) of these presumably memory-related GABAergic engram neurons, whether at recent (2 days post-conditioning) or remote time points, failed to induce similar memory-like behavior or ensemble activities (Fig. 4g,h; Extended Data Fig. 12). These results show that targeted reactivation of GABAergic USNs is sufficient for memory retrieval at the remote stage.

Do USNs gate the general retrieval of memories, or are they specific to particular memories? To investigate this, we tested if different ensembles of USNs regulate the retrieval of distinct contexts. To this end, we implanted mice with two distinct memories via successively conditioning them to two different contexts at specific time points, with 24-hour intervals (Fig.4i), since USNs were selected based on comparisons of remote activities and baseline activities occurring 24 hours apart. We were able to identify two subsets of “USNs” likely associated with the two distinct contexts respectively (Fig.4j). Targeted activation of these two subset cells successfully retrieved remote memories specific to their respective conditioning contexts (Fig.4k). These results indicate that the formations of USNs are context-specific, and targeted optogenetic activation of context-specific USNs is sufficient to retrieve remote memories of the corresponding contexts.

### USN activation recalls cortical ensembles relevant to memory retrieval

We next investigate whether the cortical ensembles evoked by photo-activation of USNs are relevant to memory retrieval. For this purpose, we compared neuronal ensembles evoked by context (virtual stimuli herein) or optical activation, and found that they were highly similar in the overall spatial pattern of ensemble activity (Fig.5a)^43,55^. In contrast, random photo-activation of the same number of memory-related GABAergic neurons (other than the USNs) only transiently activated a small number of cells (Extended Data Fig.12b) that differed from ensembles evoked by conditioning context (Extended Data Fig. 12c). The functional connections among these evoked cells, measured in pairwise neuronal synchronization counts, were also weaker than those under the contextual recall (Fig. 5b,c; Extended Data Fig.12d). We further calculated the cosine similarity of the ensemble-representing binary vectors (see Methods for details)^49,55,56^, confirming the high similarity between contextually and opto-genetically evoked ensembles (Fig.5d,e). This finding was further strengthened by the two-context experiment (Fig. 4i), showing that optical activation of different sets of USNs evoked ensembles similar to the context-specific ensembles (Fig.5f,g). Thus, direct activation of the USNs reactivates cortical ensembles relevant to the retrieval of specific remote memory.

**Fig. 5.**
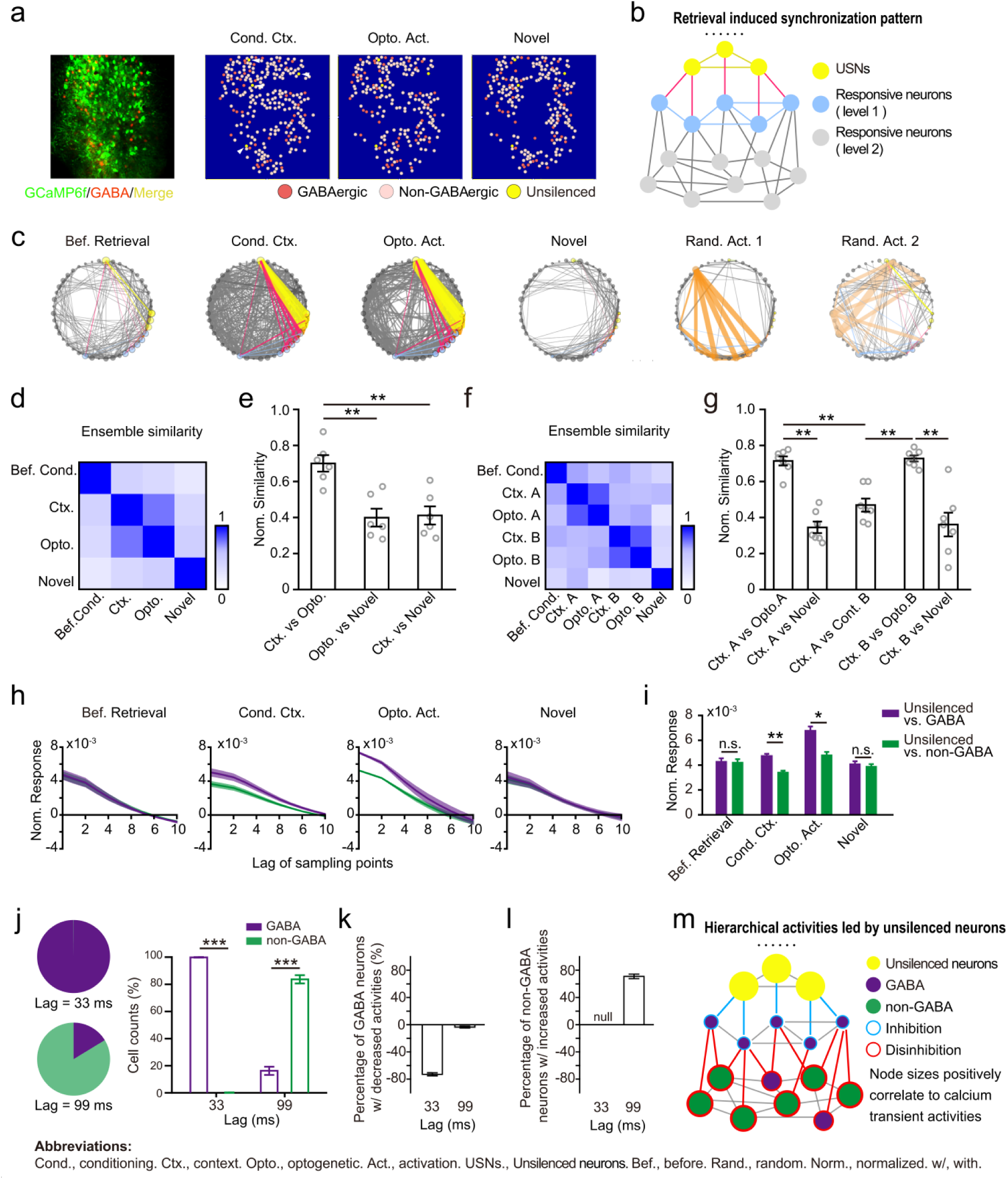
USN activation recalls hierarchical functional networks relevant to memory retrieval. **a**, Ensemble patterns (displayed by dot distribution) under contextual recall and optogenetic stimulation were similar compared to those under novel context. Left, two-photon images averaged from all time points. Right, diagrams of neurons activated upon memory retrieval under various conditions. **b**, Topological hierarchy of neuronal synchronization pattern during successful retrieval. Yellow, USNs; blue, follow-up activated neurons within 1 sampling point (33 ms) after USN activation; gray, all other neurons. **c**, Similar topological structural dynamics were observed under Cond. Ctx. and Opto. Act. As the control, neither novel context (Novel) nor random GABAergic neuron activation (Rand. Act. 1 and Rand. Act. 2) exhibited similar effects. Color semantics match B. Orange, randomly selected GABAergic neurons. **d**, Average cosine similarity of population activities under different stimuli in 1 animal. The highest ensemble similarity was found for Opto. Act. and Cond. Ctx. **e**, Normalized average cosine similarity confirmed the highest similarity between ensemble activities under Opto. Act. and Cond. Ctx. (70.66 ± 4.61, *P* < 0.01 compared to cond. and novel context similarity, *P* < 0.01, n = 6 animals, one-way ANOVA with Tukey’s post hoc test, mean ± s.e.m.). **f**, Targeted activation of the two context-specific subsets of USNs elicited distinct neuronal ensembles with similar architecture specific to their respective conditioning contexts. **g**, Normalized average cosine similarity analysis confirmed the high similarity between the neuronal ensembles elicited by Opto. Act. and their respective contexts [(Ctx. A *vs.* Opto. A) *vs*. (Ctx. A *vs*. Novel), *P* < 0.01; (Ctx. A *vs.* Opto. A) *vs.* (Ctx. A *vs.* Ctx. B), *P* = 0.0009 < 0.01]. In contrast, the similarity was low between neuronal ensembles elicited by two different contexts [(Ctx. B *vs.* Opto. B) *vs.* (Ctx. A *vs.* Ctx. B), *P* = 0.0004 < 0.01, n = 7 animals, one-way ANOVA with Tukey’s post hoc test, mean ± s.e.m.], supporting that context-specific memories are represented by distinctive ensemble patterns in the ACC. **h**, GABAergic neurons preferentially responded to USN stimulation. Lag of sampling points denotes the number of frames (33 ms per frame) following USN stimulation. Normalized responses (Y axis) calculate the normalized number of neurons (Mean ± s.e.m.; separated into GABAergic and non-GABAergic groups) that responded to USN activation. Normalization is performed by taking the ratio between each value and the mean. Color scheme is illustrated in i. **i**, Average normalized response of 10 frames (all points in h) following USN activation. Under Cond Ctx, more GABAergic neurons responded to USN activation (*P =* 0.004 *<* 0.01), likewise for Opto. Act. (*P =* 0.0104 *<* 0.05, n = 40 neurons from 5 animals, unpaired t-test, mean ± s.e.m.). **j**, The percentage distribution of GABA or non-GABAergic cells following activation of USNs. Lag = 33 ms, GABA (99.87 ± 0.13) *vs*. non-GABA (0.13 ± 0.13), P < 0.01; Lag = 99 ms, GABA (16.29 ± 3.03) *vs*. non-GABA (83.71 ± 3.03), *P* < 0.01, paired t-test. **k,l**, Most GABAergic neurons responsive to USN activation were identified in the sampling lag within 33 ms and displayed decreased activity (k). In contrast, non-GABAergic neurons with increased activity were largely observed at 99 ms sampling lag (l). **m**, Framework of the activation pattern under successful memory retrieval. Activation of non-GABAergic neurons (mostly in 99 ms) lagged behind GABAergic neurons, suggesting a disinhibitory network downstream of USNs.

How can photo-activation of a small number of GABAergic inhibitory neurons evoke entire neuronal ensembles relevant to memory retrieval? One possibility is that the evoked neuronal ensemble constituted a small-world architecture under the control of GABAergic USNs for efficient communication^57–62^. In this model, GABAergic USNs act as hubs, synchronously amplifying input signals by inhibiting other non-unsilenced GABAergic neurons^63^. To identify the excitatory/inhibitory nature of the hierarchical responses, we examined neurons that responded within 10 imaging frames (330 ms) following the activation of USNs (Fig. 5h, i). We found that most GABAergic neuronal responses were identified within one imaging frame (33 ms) following activation of USNs during memory retrieval. Notably, these responsive GABAergic neurons displayed decreased activity (Fig. 5j). In contrast, non-GABAergic neurons with increased activity were primarily observed at 99 ms sampling lag (Fig. 5k). This suggests that targeted activation of USN triggers downstream non-GABAergic neurons through a disinhibitory mechanism^64^. We found further support for this hypothesis via community analyses for ensemble activities under different retrieval conditions (Extended Data Fig. 13)^65^. Our results revealed that coactive firings of USNs and the subsequent ensemble activation constituted hierarchical layers of modules. Notably, with USNs forming the first module layer, second module layer neurons were entirely GABAergic, confirming analyses in Fig. 5j. Furthermore, the neuronal ensembles triggered by optogenetic stimuli showed similar modularity-hierarchy distributions to those triggered by conditioning context (Extended Data Fig. 13a). However, the modularity distribution evoked by novel context differed significantly (Extended Data Fig. 13d). These results confirm similarity analyses in Fig. 5d to 5g and indicate that the functional modularity structures elicited upon USN activation were crucial for successful memory retrieval. Collectively, USN activation recalls remote memory via an indirect disinhibitory activation of specific ensembles.

### Unsilenced neuronal ensemble gates remote memory retrieval

To further investigate whether the GABAergic USNs were required for gating remote memory retrieval, we photo-suppressed them while the animals were exposed to the conditioning context on post-conditioning day 63. We co-expressed GCaMP6f and the light-driven chloride pump halorhodopsin eNpHR3.0 in the ACC region and conducted photo-suppression on the eNpHR3.0+ USNs (Fig. 6a,d). Our initial attempt to image and photo-inhibit USNs in one imaging plane (2D imaging and photo-stimulation) only elicited a minor tendency towards suppression of remote memory retrieval, which was not statistically significant (Fig. 6b). We speculate that contextual cue-triggered remote memory retrieval might engage activation of more USNs scattered at different depths of the ACC region. Accordingly, inhibiting USNs in the imaged ACC volume may be required for behavioral suppression.

**Fig. 6.**
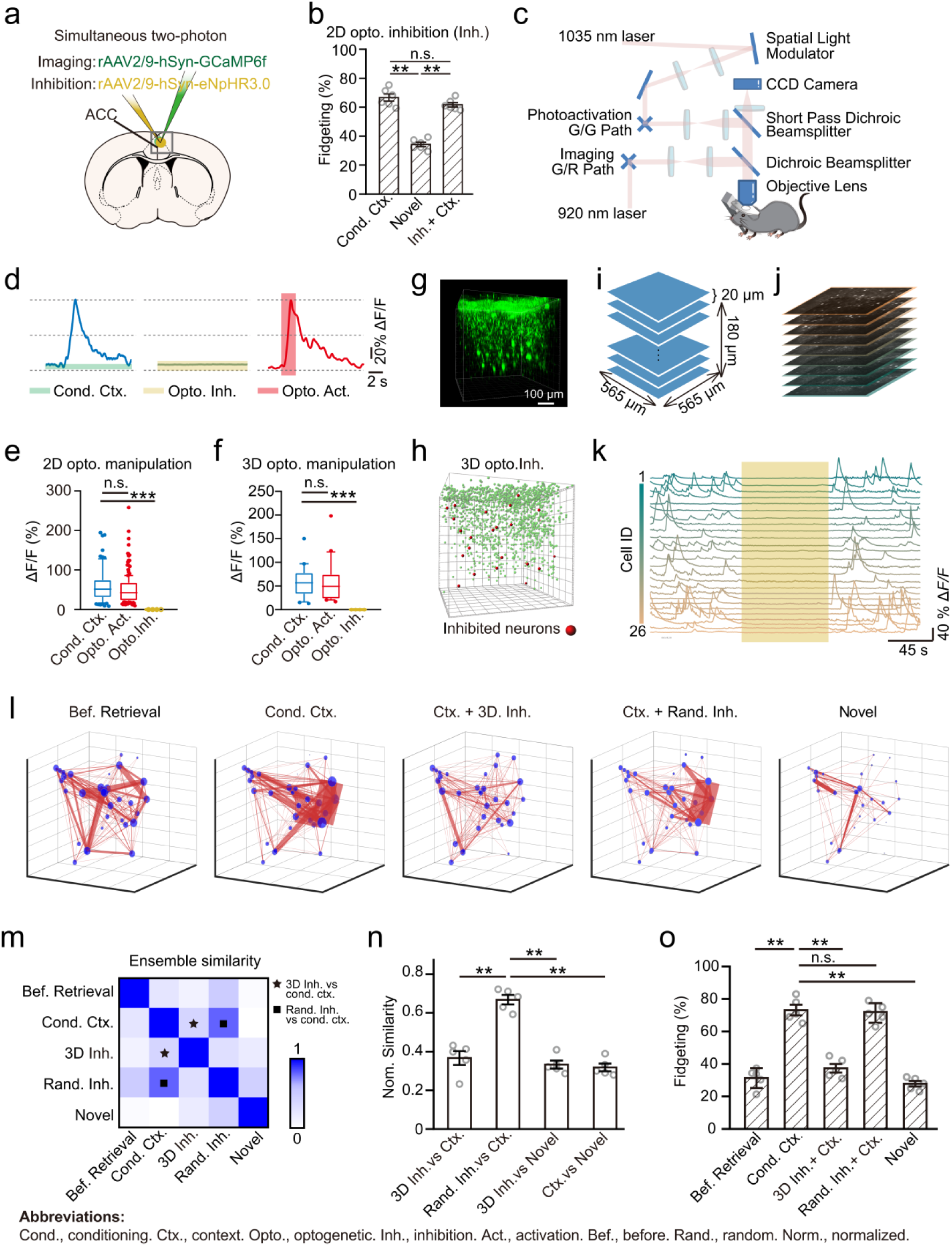
Targeted 3D holographic suppression of USNs impairs remote memory retrieval. **a**, Viral injection schematics. **b**, 2D holographic optogenetic inhibition failed to inhibit the retrieval of associative memory. **c**, Schematics of microscope setup. Two ultrafast lasers combined with galvo/resonant (G/R) and galvo/galvo (G/G) paths were employed respectively for imaging and spatial light modulator stimulation. **d**, Raw calcium ΔF/F response curve of an example USN show effective optogenetic inhibition. **e**, All photo-stimulated USNs were effectively inhibited during the 2D manipulation (Cond. Ctx. = 60.30 ± 5.01 %, Opto. Act. = 51.09 ± 2.76 %, Opto. Inh. = (5.39 ± 2.63) × 10^-4^ %, Cond. Ctx. *vs* Opto. Act, *P* = 0.18, Cond. ctx. *vs* Opto. Inh., \*\*\**P <* 0.001, one-way ANOVA with Tukey’s post hoc test, n = 6 animals, mean ± s.e.m.). **f**, Photo-stimulated USNs were effectively inhibited during 3D manipulation. (Cond. Ctx. = 58.15 ± 7.35 %, Opto. Act. = 57.97 ± 9.51 %, Opto. Inh. = (5.38 ± 2.62) × 10^-4^ %, Cond. Ctx. *vs* Opto. Act, *P* = 0.18, Cond. Ctx. *vs* Opto. Inh., \*\*\**P <* 0.001, one-way ANOVA with Tukey’s post hoc test, n = 5 animals, mean ± s.e.m.) **g,h**, 3D reconstruction diagram of the entire imaged neuronal population in a volume of 565×565×180 μm^3^. n = 867 neurons in 1 animal. Inhibited USNs are marked in red in H (n = 26 neurons). **i,** 3D volumetric imaging was achieved via piezoelectric module-controlled lens movement. Each imaging plane was spaced 20 μm apart with 250 ms total collection time for each volume. **j,** Example 3D image stacks. **k**, Effective complete inhibition of all USNs in 3D space shown by ΔF/F curves in a sample animal (n = 26 neurons). **l**, 3D coactivity network patterns demonstrate suppression of pairwise neuronal synchronization by volumetric inhibition of USNs. Note that for better visualization, only USNs are shown. **m**, Cosine similarity analysis revealed that targeted 3D inhibition of USNs decreased the ensemble similarity to that elicited by Cond. Ctx. **n**, Normalized average cosine similarity confirms lower similarity in 3D Opto. inhibition (Opto. Inh.) *vs* Cond. Ctx. compared to Rand. Inh./Cond. Ctx. **o**, Successful suppression of remote memory retrieval via volumetric targeted inactivation of USNs. Under the conditioning context, inhibition of USNs reduced the fidgeting rate to a level comparable to novel context control (Cond. Ctx. *vs*. Bef. Retrieval, *P* < 0.01, Cond. Ctx. + 3D Inh. vs. Bef. Retrieval, *P =* 0.454 > 0.05). In contrast, inhibition of randomly selected GABAergic cells did not reduce fidgeting (Cond. Ctx. + Rand. Inh. *vs*. Bef. Retrieval, *P* < 0.01, Cond. Ctx. +3D Inh. *vs.* Cond. Ctx., *P* < 0.01, one-way ANOVA with Tukey’s post hoc test, n = 5 animals, mean ± s.e.m.).

To address this, we collaborated with microscopy engineers (Thorlabs, USA) to achieve simultaneous volumetric imaging and optogenetic manipulation of USNs using double light paths and a 3D-adapted spatial light modulator (SLM) (Fig. 6c)^66^. Inhibitory effects were again confirmed by fluorescence trace response and average activities of all the targeted USNs (Fig. 6d-f). To improve imaging coverage of the ACC region, we performed longitudinal 3D imaging at a volume of 565 × 565 × 180 μm^3^ at the ACC (Fig. 6g-h; see details in Methods). We identified much more (28-38 neurons) GABAergic USNs spanning multiple (10) imaging planes (Fig. 6h). Photo-stimulation of eNpHR3.0+ USNs flattened their calcium response, dampened the coactivity levels (Fig. 6k,l), induced 3D neuronal ensembles that were different from those elicited by the conditioning context or a novel context (Fig. 6m,n), and dramatically suppressed remote fear memory retrieval (Fig. 6o). As a control, volumetric optogenetic inhibition of randomly selected neurons failed to elicit the above effects (Fig. 6l-o). These results highlight the necessity of suppressing a sufficient number of GABAergic USNs to prevent memory retrieval. Together with the USN photo-activation results, our findings collectively indicate that unsilenced GABAergic neurons reliably gate remote memory retrieval.

## Discussion

Uncovering the trace for remote memory has always been a challenge in the field, as the memory traces activated during encoding subsequently undergo reorganization during consolidation and ultimately persist in a dormant state. To retain their potential for being retrieved memory traces may possess “retrieval handles” that permit reactivation by specific representations^7,17,21,22,67^. Here, we reveal at the remote memory stage a small number of cortical GABAergic neurons that displayed functional features of “retrieval handles”. These USNs displayed modest activity “silence” after conditioning, but their activities never completely decreased to the before-conditioning baseline level. Instead, they maintained their increased calcium power and ensemble connectivity at a modest above-baseline level throughout the remote stage. The connectivity may function as a specialized “tag” for memory retrieval. Because of the elevated and synchronized activity, the tagged trace could be easily tracked down upon memory retrieval and thus may be employed as the “retrieval handle”. Our results with bidirectional targeted manipulations of the activity of these cortical neurons confirmed their causal role in gating the retrieval of remote memory. The default intermediate activity level of these USNs remained substantially lower than that at the conditioning or retrieval stage, explaining why they would not recall the memory constantly.

The cortical neurons we uncovered here possess at least four characteristics. Firstly, they exhibited persistently higher activity than other neurons entering the dormant state. Despite a significant activity decrease since the memory encoding stage, they maintain an intermediate activity level above the before-conditioning baseline throughout the entire long-term memory stage. Secondly, the number of USNs is stably unchanged. Thirdly, they displayed a higher degree of inter-neuronal synchronization. The synchronous activity within these neurons implies the potential formation of a cohesive control unit that collaborates to perform retrieving functions. Fourthly, they were all identified as GABAergic neurons. Despite the limited population of these neurons, their activation as inhibitory interneurons enabled them to exert control over a larger number of downstream neurons.

These characteristics were partially in line with memory engram ensembles discovered in the past. Various studies have found functionally distinct subsets of cells with unique molecular markers that specify contextual information among engram ensembles^60,61,68–71^. However, the few numbers of USN and unique coactive features still set them apart from previously uncovered engram subsets. This leads us to wonder what is the relationship between the USNs and the conventional memory engram cells. There remains the possibility that engram cells themselves have tight networks and activation of engram subpopulations not just specific to USNs can also activate entire memory-related network. To address these questions, we compared stimulation of USNs and stimulation of random memory-related GABAergic cells activated at the conditioning stage. Referring to the engram cell tagging technique^23,60,61,68^, we identified GABAergic cells with increased activities during the conditioning stage and then examined the effect of photo-stimulation of the same number of random GABAergic neurons. Furthermore, this stimulation failed to elicit functionally meaningful downstream patterns found in successful memory retrieval induced by either natural context or stimulation of the USNs. These results suggest that the retrieval of memory-like behavior is not simply elicited by random reactivation of engram subpopulation. Instead, targeted reactivation of GABAergic cell subpopulation specific to USNs gates memory retrieval at the remote stage, possibly via triggering downstream hierarchical network.

The downstream hierarchical network led by USNs could serve as a "cascaded amplification" mechanism to effectively recall downstream neurons, enabling cortical representations of remote memories upon retrieval^49,72–74^. This phenomenon helps explain why we were able to observe behavioral responses by just stimulating a few cortical interneurons. The contextual cue might activate USNs scattered at different imaging planes upon memory retrieval. In line with this speculation, we found that volumetric suppression of USNs ensemble impaired memory retrieval. This highlights the USNs’ function as “retrieval handles” to gate larger engram populations downstream^22,28,29,75–78^. The “gating” function can be realized by a local disinhibitory circuit, as suggested by our framewise analyses of cellular components following USN activation^59,64,76,79,80^. In addition, as the ACC serves as a primary associative cortical area integrating sensory and hippocampal inputs, sparse neuronal codes from upstream sensory areas could activate USNs^28,29,51^. The USNs may thus integrate upstream hippocampal inputs and gate downstream remote memory traces likely via the disinhibitory multilayered network, maintaining their dormancy until being reactivated by corresponding sensory cues^11,12,22,28,64,80^.

Functional division and organization principles of memory trace post-system consolidation have emerged as one of the most important focuses within the field^5,48,59,60,62,74^. Here we reveal for the first time a small cortical inhibitory ensemble that serves as theorized “retrieval handles” to gate the retrieval of a two-month memory in mice. The unsilenced neuronal ensembles were strictly dissociated based on persistently heightened and coactive features throughout the remote stage. More notably, conditioning the mice to two different contexts subsequently formed two distinct clusters of USNs that can be selectively photo-activated to retrieve respective memories. Future work could either aim to identify established GABAergic subtypes of USNs or identify new molecular markers via single-cell sequencing techniques^59,64,80^. The current results support functional connections between USNs and downstream neuronal networks they control, which likely include latent memory traces. By using more sensitive probes with higher spatial-temporal resolution, we would be able to examine the universalities of the gating mechanisms and further uncover the organization principles of the downstream memory trace in the mammalian brain.

## Acknowledgements

We thank Dr. Lu-Yang Wang, Dr. Peng Yuan and all members of the Lu lab for discussion and support.

## Funding

The Science Technology Innovation 2030 Project of China (2021ZD0203502 to WL)

The National Natural Science Foundation of China (T2394531 to WL)

The Ministry of Science and Technology of China (2021YFA1101302 to WL)

The National Natural Science Foundation of China (32200835 to SLW)

The China Postdoctoral Science Foundation (2021M700847 to SLW)

The China Postdoctoral Science Foundation (2024T170168 to SLW).

## Author contributions

Investigation: SW, TS, FS, HY, RC

Supervision: WL, CZ

Funding Acquisition: WL

Visualization: SW, TS

Writing: WL, SW, HY

## Inclusion and diversity

We support inclusive, diverse, and equitable conduct of research.

## Resource availability

### Lead contact

Further information and requests for reagents and resources should be directed to W. Lu.

## Material availability

This study did not generate new unique reagents.

## Data and code availability

- All data and analyses necessary to understand and assess the conclusions of the manuscript are presented in the main text and in the supplementary materials.
- All the analysis codes are available from the lead contact upon request.

## Supplementary information

### Methods

#### Animals

All animal procedures were performed under the guidelines of the Institutional Animal Care and Use Committee of Fudan University. Wild-type C57BL/6 male mice were purchased from GemPharmatech (Nanjing, China) and Vital River Laboratories (Beijing, China). To distinguish GABAergic and non-GABAergic neurons, *Vigat*-cre (Jackson Laboratory, strain No: 017535) were crossed with Ai9 mice (Jackson Laboratory, strain No: 007909) to obtain GABA^+/Cre^;Ai9^wt/mut^ (GABA-Cre::Ai9) mice. Genotypes were determined via PCR postnatally at 21 days. All mice were group housed in five at the SPF facility of the Department of Laboratory Animal Science at Fudan University. Animals were maintained on a 12-hour reverse light-dark cycle with *ad libitum* access to water and a chow diet. All experiments were performed during the light cycle of mouse subjects. All mice were aged 2-5 months old at the time of viral injection. The number of subjects in each treatment group was determined based on expected variance and effect size from previous imaging and behavioral studies.

#### Stereotactic and head plate attachment procedure

Before surgery, the mice (4 weeks of age) were anesthetized with tribromoethanol by intraperitoneal (i.p.) injection (240mg/kg, Sigma-Aldrich, USA). Local anesthesia and post-operational pain management were given by lidocaine hydrochloride gel (Xinya Med Co., China) applied to the skin around the incision. A thin layer of erythromycin eye ointment (Baijingyu Med Co., China) was applied to the eyes for protection. Body temperature was maintained at 37 ℃ with a small heating pad. The target surface of the mouse skull was exposed with a ring incision after shaving the head. The periosteum over the exposed skull surface was removed using a scalpel and hydrogen peroxide (3%). The location of ACC region (AP 0.26 mm; ML -0.35 mm; DV 0.3 mm) was then determined via the standard stereotactic procedure and a cranial window (5×5 mm^2^ size) was opened directly above the targeted region followed by viral injections.

For optogenetic stimulation, a cocktail of viruses rAAV2/9-hsyn-GCaMP6f-WPRE (500 nl, 2e12 vg/ml) and rAAV2/9-hsyn-C1V1(T/T)-TS-P2A-WPRE (500 nl, 2e12 vg/ml) were injected into ACC region (AP 0.26 mm; ML -0.35 mm; DV 0.3 mm) using pulled glass pipettes. For optogenetic inhibition, C1V1 was replaced for rAAV2/9-hsyn-eNpHR3.0-(T/T)-TS-P2A-WPRE (500 nl, 2e12 vg/ml). The injection rate was set at 30 nL/min, and the pipette was held for 15 min post injection to avoid fluid flow back due to pipette removal. A 5-mm circular glass coverslip was placed and sealed by cyanoacrylate adhesive for chronic imaging.

After the cranial window was closed by the glass coverslip, a custom-designed titanium head plate was attached to the skull by dental cement (Yuyan Instruments, China). Postsurgical dexamethasone sodium phosphate (2 mg/kg, Shanxi Ruicheng Kelong Veterinary Medicine Co., Ltd. China) and ceftriaxone sodium (200 mg/kg, Guangxi Kelun Pharmaceutical Co., Ltd. China) were respectively injected subcutaneously and intraperitoneally daily for a week. During this period mice recuperated in the home cage with food and water *ad libitum*. Ceftriaxone sodium was also injected daily for 12 weeks during the entire experiment to prevent inflammation.

#### Two-photon imaging protocol

The imaging system employed in our study was Thorlabs’ Bergamo® II Series Multiphoton Microscope (Thorlabs, USA). For two-photon imaging, the Coherent Chameleon Vision II (Coherent Corp. USA) Ti:Sapphire Laser (920 nm) was employed as the laser source coupled with an 8kHz Galvo/Resonant (G/R) scanner. For optogenetic stimulation and inhibition, the Coherent Monaco 1035 nm femtosecond laser was used to excite red-shift opsins C1V1 or eNpHR3.0, coupled with a Galvo/Galvo (G/G) scanner. Simultaneous imaging and holographic optogenetic manipulation were thus achieved respectively via the G/R and the G/G path. The laser power, stimulation positions, and patterns were quality-controlled by their corresponding commercial software (ThorImage 4.1, USA).

At each data acquisition time point, mice were awake and head-fixed on the platform. To realize effective longitudinal tracking of the same group of neurons, reference brightfield images of the cortical surface and two-photon images of the targeted area were acquired for each animal during “before-training” baseline recordings. In all subsequent imaging, hardware coordinates were checked and representative features (blood vessels and each neuron) within the imaging window were manually re-aligned with the reference image. Images were acquired at 30 Hz for 90s (2800 frames) at each imaging time point. 16 × objective, 0.8 NA, Nikon object lens was employed. The field of view (FOV) was set at 512 × 512 pixels (565 × 565 μm^2^). The laser power was strictly monitored for an average value smaller than 80 mW across all sessions to reduce phototoxicity and bleaching.

3D volumetric imaging was achieved via fast-scan piezoelectric modules. The module controlled the movement of the objective lens in a 400 μm range via voltage-dependent deformation. During 3D imaging, the focal plane was moved vertically at 20 μm per step upon complete acquisition of one 2D imaging plane (Fig. 6i). The 20 μm step length was determined based on the average soma size of cortical neurons (19.1 μm) to minimize inter-neuronal signal interference along the z-axis. The average soma size was measured from 400 randomly selected neurons in 2 animals under 3D scan. Each imaging plane consisting of 512×512 pixels (565 × 565 μm^2^) was acquired via the Galvo/Resonant scanner in a raster pattern, moving in the two-way scan mode. A total of 10 planes were obtained to construct a 3D FOV of 565 × 565 × 180 μm^3^. For each animal, the deepest imaging plane with clearly visible neurons was selected as the sampling start point. The sampling starting depth ranged from 560-500 μm and the ending depth ranged from 380-320 μm below the pia surface. Each imaging plane costed 33 ms (at 30 Hz sampling rate) to complete. The effective sampling rate was 3 Hz. The time dwelt on each 3D field of view (FOV) was 330 ms.

As ACC is mostly covered by neighboring motor cortex, we took two measures in ACC targeted imaging experiments to corroborate the region-specificity. First, we referred to the previous study and employed the middle artery as a reference for positioning.^38^ Second, we replicated the exact same imaging experiments in the motor cortex (AP:0.26 mm, ML:-0.7 mm, DV:0.3 mm) lateralized to targeted ACC area. Based on the quantitative definition for USNs, we did not detect any USNs in neighboring cortical regions, suggesting no USNs could be identified in the motor cortex. The USNs we observed were thus indeed located in surface ACC region.

#### Holographic optogenetics protocol

Holographic imaging and stimulation were conducted using the commercial Bergmo II microscope (Thorlabs, USA) installed with a volumetric spatial light modulator (SLM) module. In-vivo holographic stimulation was first performed sequentially as quality control and then simultaneously for the formal experiment. Sequential stimulation mode employed the spiral scan feature of the Galvo/Galvo scanner (A1R MP+, Nikon). Simultaneous holographic stimulation was achieved via the SLM module established in collaboration with Thorlabs. The light path was established based on designs by Häusser and Yuste’s labs. ^50–52,66,81^ and more detailed information could be obtained from Thorlabs (https://www.thorlabs.com/) upon request. Two laser oscillators were integrated respectively to the imaging (Coherent Chameleon Vision II with 80 MHz pulse freq., 140fs pulse width used at 920 nm) and stimulation (Coherent Monaco with 80 MHz pulse freq., < 400fs pulse width fixed at 1035 nm) light paths of the Thorlabs Bergmo II microscope. The SLM window was also separated into two regions for simultaneous modulation of imaging and stimulation paths lasers. For beam width registration of the two imaging and stimulation lasers to their assigned SLM regions beam-expanding telescopes were employed following the pockels cells. To rule out interference from the two regions we employed spatial filters (SF) in conjugated pupil planes and a zero-order block (ZOB) for removal of zero-order beam. The entire holographic control volume approximately covered 600 × 600 × 400 μm^3^.

The diameter of stimulation ROIs was set to be consistent with the observed averaged cell body size. The laser power was set at 5-20 mW on each neuron. The specific power of the optical stimulation was analyzed and examined via commercial software (ThorLabs, USA). Activation effects for both sequential and SLM stimulation were tested first on a fluorescent plate and then in vivo (Extended Data Fig. 11). Pre-stimulation SLM calibration was performed via the commercial coupled software (ThorLabs). The calibration software automatically computes the transformation between SLM target coordinates and imaging FOV coordinates on the burnt locations on the fluorescence plate using methods described previously.

For results reported in Fig. 4, all identified co-active neurons were simultaneously stimulated using five sweeps via SLM modules (Thorlab, USA). Stimulation frequency is 4Hz and each sweep lasts 2 seconds. Animals (n = 3) that did not show full response (ΔF/F > 3σ) to all sweeps were excluded. In the remaining 12 animals, 10 animals successfully retrieved remote memories upon target activation of USNs. The stimulation success rate of 100 % per sweep and average response of 0.69 ± 0.15 (norm. ΔF/F) is comparable to previous literature.^52^

For the random stimulation results reported in Fig. 4 and Extended Data Fig. 12, stimulation targets were selected in the following steps. First, all identified GABAergic neurons in the imaging plane were given a number ID. Then a fixed number of IDs were randomly selected as stimulation targets. The number of IDs selected was determined by the number of identified USNs in that subject (for specific numbers please see supplementary tables).

All optogenetic manipulations were delivered on day 63 thirty minutes post memory retrieval tasks. To assess the causal role of USNs in memory retrieval, a total 3 min of 3D optogenetic inhibition was delivered on selected USNs accompanied by training context (context A) presentation. Inhibition was separated into two trials (1.5 min each, with 2 min ITI within) to limit thermal damage on the neurons.

#### Behavioral assays

The behavioral paradigm was adapted from the classical contextual fear conditioning task which consists of a training block under context A and day 63 retrieval tasks under context A, a novel context B and optogenetic manipulation-induced artificial retrieval. We used a mini-computer (Raspberry Pi) controlled tablet screen to display the virtual context to the animal at the start of each trial. During training trials, the screen displayed a wall corner stacked with geometric square elements; during novel context trials, the screen displayed an open room stacked with real-life elements (e.g. furniture and sofas). For optogenetic stimulation, the screen was turned off, and the animal was kept in a default dark environment.

A week post viral injection, each animal was placed on a head-restraint apparatus under the two-photon microscope for 30 -60 min daily (for a total of 21 days) to reduce restraint brought-on anxiety and stress and adapt to the experimental environment during the imaging process.^82^ On the first day, the mice were exposed to the training environment for 3 min without any stimulations before the conditioning training. Then the virtual context (conditioned stimulation, CS) was delivered and the tone stimuli were set at 2 kHz, 80 dB for 20 s by an audio controller. Nociceptive stimuli (unconditioned stimulation, US) were delivered via the DM-300 CO_2_ laser stimulator (20 ms, 8 W; custom designed by Dimei Optics & Electronics Tech. Co. Changchun, Jilin, China) attached to the animal’s back paw. The training session contained six consecutive training blocks with an ITI of 40 s, each training block consists of a 20 s baseline, a 20 s, 80 dB tone, followed by an 18 s trace interval of silence and a 20 ms, 8 W nociceptive foot shock. On the retrieval day, mice were re-exposed to the conditioning context to assess contextual conditioned fidgeting behavior for 3 min. Then after 30 min rest, mice were exposed to a novel virtual context for 3 min as the control. 3 min after the onset of novel context presentation, mice were also presented with the 20s tone CS, and demonstrated increased fidgeting during tone presentation (53.2% vs. 29.2% before tone CS presentation, please see Extended Data Fig. 14a).

For the two-context test, each animal first underwent the entire training session under context A and then 24 h later under context B. On the retrieval day, mice were re-exposed to context A and 24h later to context B. On retrieval test day 3, mice were presented with a novel context C to assess the contextual specificity of the associative memory. The novel context C consists of two rooms, one furnished with gray floors, blue walls, and pink sofas and one furnished with wooden floors, green walls, and green sofas.

Control experiments were performed to test if the formation of USNs relies on tone presentation. Animals were conditioned without tone (spatial context+ nociceptive laser) and presented the conditioning context during retrieval sessions. On the training day, animals were exposed successively to context A and context B. The order in which we present the two contexts was counterbalanced across animals. Context A consists of square building blocks with metal floors. Context B consists of triangular building blocks with coarse cardboard floors. For each trial, a 2 s, 1mA foot shock was administered 38s upon context presentation. 6 trials were repeated with an intertrial interval of 30 s. On the retrieval day, context A, context B, and a novel context were successively presented, each for 3 min. The order of presentation was again counterbalanced across animals. The novel context consists of irregular quadrilateral building blocks with metal floors. Our results demonstrated that animals successfully formed memories respectively for context A and context B (Extended Data Fig. 14b). The behavioral performance during 63 d memory tests, however, was inevitably weakened compared to training with tone. Collectively, our results show that the formation of USNs and the retrieval of remote memory is independent on tone stimuli.

To test if the formation of USNs were influenced by the head-fixed paradigm, we also replicated the entire assay in standard fear conditioning box. Animals were first habituated to the imaging platform for 21 days and then trained in a classical fear conditioning box. Fear conditioning was carried out in the default background (grid floor, plexiglass walls), consisting of 6 trials of 20 s, 8 kHz tone, 18 s trace delay, and 2 s, 1 mA foot shocks. The intertrial interval was 30 s. After training, animals were imaged at selective time points (30 min, 28 d - 63 d). Mice successfully formed associative memories under this paradigm (Extended Data Fig. 14b), with an overall decrease in freezing compared to fidgeting on the imaging platform. Our new data demonstrated that USNs can still be identified under the free-moving training paradigm, indicating that they are not a phenomenon specific to the head-fixed fidgeting behavioral assays.

Experimental designs are illustrated in Figures 1a to c. The movement of the animal’s forepaws was recorded as a change in the continuous voltage signals being recorded throughout the entire experiment. All the virtual context cues, sound cues, pain delivery, fidgeting movements, and imaging acquisition were controlled by the EPC-10 amplifier (Heka, Germany) to synchronize delivery and recording.

#### Fidgeting rate analysis

The fidgeting rate was measured from voltage changes in a piezoelectric sensor positioned under the animal’s forepaws. Absolute value of raw signals from the piezoelectric sensor was plotted in Fig. 1h. For fidgeting behavior analysis^33,34^, the recorded continuous voltage signal was down-sampled to 100 Hz, and then the average voltage throughout the entire recording time was calculated as F0. Relative changes in signals were calculated by (F-F0)/F0. To eliminate negative values, the amount of fidgeting was calculated by taking the root mean square of the voltage signal. To quantify behavior associated with conditioning, the average fidgeting rate 3 min before training was calculated as the baseline, and the percent time of fidgeting movement above 3 standard deviations (SD) from the baseline was calculated for each experiment session.

To validate the results on fidgeting behavior, we employed video analysis to assess the specificity of the sensor for picked "fidgeting" behavior. Specifically, we trained the animal under head-fixed conditions in a conditioning box under the exact same protocol used in previous experiments. Then we video-recorded the entire behavioral assay on the imaging platform and analyzed the grip and release motion of the animal’s front paw. Each grip of the animal’s front paw was labeled as a “fidgeting” response via frame-to-frame comparison analysis, which we termed immobility. The percentage of immobility calculated the degree of motion (frame-to-frame difference) in the video and small immobility values correspond to front paw gripping (similar to “freezing” in classical fear conditioning video analysis). The gripping actions increased front paw pressure, as captured by increases in raw piezoelectric signals. Therefore, small immobility values correspond to animals’ conditioned fidgeting response, whereas large raw piezoelectric signals correspond to fidgeting behavior (Extended Data Fig. 1a). The exact inverse relationship between immobility and piezoelectric trace shown in Figure 1a demonstrated a good match in results from these two methods. We calculated the matching rate of immobility responses and the piezo-electric recorded signals as 77.56% (Extended Data Fig. 1c). From this value, we obtained a mismatching rate of 22.44% (100%-77.56%). To further assess whether the 22.44% mismatching rate would cause bias in behavioral tests, we generated 1000 simulated fidgeting responses at baseline and remote stages (in a range of true values ± 22.44%). We conducted t-tests using these simulated values and found that 99.6% of p-values were significant (<0.05, Extended Data Fig. 1d,e). Since the original t-test based on true fidgeting values was also significant (<0.05), this result indicated that the conclusion of animals significantly responding to conditioned stimuli during remote retrieval remained the same no matter for simulated or true results. The small chances of mismatch between piezoelectric signals and immobility could not override the conclusion on remote memory retrieval. Together these results suggest that the signals-labeled “fidgeting” response could be used as a reliable readout for the behavioral expression of memory.

To further examine if other behavioral changes that would corroborate our findings with fidgeting, we employed the fear conditioning protocol (e.g. freely moving tone fear conditioning followed by 2P imaging) to see if they can elicit activation of ACC excitatory neurons. Animals were first habituated to the imaging platform for 21 days and then trained in a classical fear conditioning box. Fear conditioning was carried out in the default background (grid floor, plexiglass walls), consisting of 6 trials of 20 s, 8 kHz tone, 18 s trace delay, and 2 s, 1 mA foot shocks. The intertrial interval was 30 s. After training, animals were imaged at selective time points (30 min, 28 d -63 d). Mice successfully formed associative memories under this paradigm (Extended Data Fig. 1f), with an overall decrease in freezing compared to fidgeting on the imaging platform. Our preliminary results demonstrated that USNs can still be identified under free-moving training paradigm, indicating that they are not a phenomenon specific to the head-fixed fidgeting behavioral assays. Together, these results suggest that the signals-labeled “fidgeting” response could be used as a reliable readout for the behavioral expression of memory.

#### Image data processing

Image processing was performed with ImageJ (National Institutes of Health, Bethesda, MD, USA) and MATLAB (Mathworks, Natick, Massachusetts, USA). To correct motion artifacts and day-to-day positional variance, we employed a calibration program using image triangulation and piecewise affine transformation (TPAT), developed by our collaborators^83^. All corrected images were also manually checked following a visual comparison method as quality control^84^. Only data presenting consistent features including cells and blood vessels within the same field of view are deemed suitable for further analysis. Any data exhibiting significant uncorrectable vibration was classified as experimental failures and is therefore excluded from statistical analyses. Regions of interest (ROIs) were manually selected using a custom ImageJ GUI that displays both the radius, location and corresponding traces of the neuron circled ^34,44^. ROIs that fully encapsulate the contour of a single neuron and have activity during the 90s imaging timeline (with typical calcium transients) were kept. To best reduce signal interference from overlapping neurons on the same imaging plane, we (1) reduced the literal size of the ROIs at an average of 15 μm to avoid potential signal overlap from a neighboring neuron or (2) rejected ROIs that are over 0.75 in Pearson correlation coefficient of signals. Approximately 10% of ROIs were rejected post-screening and cross-day comparisons (only ROIs at the same location were kept to ensure same neuron were tracked across days). Raw calcium fluorescence values were then extracted by calculating pixel grayscale intensity of the ROIs. For each ROI (ΔF/F_0_) was computed (results for USNs reported in Fig. 1h), where ΔF = (F_i_ – F_0_)/F_0_; F_0_ was calculated by the average of a 20 frames time window for a period without observant calcium events (flat baseline). For each cell, the total calcium power for each imaging time point was calculated by taking the integrals of the raw calcium fluorescence values. To compare changes in neuronal ensemble activities after contextual fear conditioning training, we calculated the calcium dynamics in each ROI across various imaging time points (before training and delays respectively at 30 min, 28 d, 35 d, 39 d, 42 d, 46 d, 49 d, 52 d, 56 d, 59 d, 63 d).

#### Quantitative definition of “unsilenced cells” (USNs)

To assess neuronal activation during the remote memory stage (day 28 to day 63, 4-9 weeks post encoding), we calculated normalized average calcium power for imaging time points at 28d, 39d, 42 d, 49 d, 52 d, 59 d and 63 d. Each imaging timepoint’s average calcium power is calculated by taking the integrals of the raw calcium trace and then dividing it over the 90 s total time. Normalization is done by subtracting the before-training’s average calcium power from each imaging timepoint’s average calcium power. Before training average calcium power was thus calculated as zero. Neurons that maintained above zero normalized calcium power for every imaging time point from day 28 to day 63 (4-9 weeks, covering the remote memory stage) are defined as USNs.

#### Corroboration of stable, training-specific feature of USNs

To determine whether the identification of USNs is influenced by arbitrariness in the selection of imaging time points, we performed random simulation experiments based on multiple remote-stage imaging time points. We randomly selected 1 up to 29 imaging time points during the remote stage from actual samples in one animal and calculated the average normalized calcium power. We then calculated the proportion of neurons with power values persistently above zero. We found that this proportion remains relatively unchanged when the total number of imaging time points considered exceeds 7 (reported in Fig. 1n). We thus selected 28 d, 39 d, 45 d, 49 d, 52 d, 59 d, 63 d, a total of 7 imaging time points one week apart to be considered in the identification process of “unsilenced cells”.

To assess whether “USNs” only existed in trained animals (CS-US group), we investigated temporal changes in average calcium power and spike frequency in CS-US and CS-only group (Extended Data Fig. 5d). Neurons with persistently elevated neuronal activities during every remote stage time points can only be observed in CS-US group, indicating that “unsilenced” neuron is training-specific. To statistically prove this observation, we also employed the bootstrapping method to compare CS-US group and CS-only group. Firstly, we constructed distribution of numbers of neurons fulfilling the quantitative definition of “USNs” from randomized results (blue lines in Extended Data Fig. 5f). The randomized results were generated by shuffling the calcium power and spike frequencies of different neurons across all imaging timepoints, making the neuronal activities time independent. We then calculated the average probability value of finding an “USN” in 1000 randomly shuffled imaging results. This averaged probability was named as “true”, though it must be noted that “true” do not refer to actual imaging observations. “True” represented the probability of neurons meeting “unsilenced” definition criteria after all data were rearranged in random scrambles. The bootstrapping results demonstrated that the “true” fell right within the resampled distribution curve in CS-US group, but not in CS-only. These analyses collectively show that USNs were unique to CS-US group.

#### Spike inference and synchronization calculation

To evaluate activation synchrony between neurons, calcium spikes representing neuronal action potentials (AP) were extracted. The time series of spike intensities for each neuron was thus converted into a binary sequence of 1 and 0 for each frame, with 1 denoting a spike and 0 denoting no spike. Since spike inference has inherent limitations and heavy dependence on the signal-to-noise ratio in the raw data, we utilized both DeepSpike inference^40^ and a comparatively more conservative deconvolution method^44^. Due to the different detection criteria in these two methods, DeepSpike has a higher inference accuracy in events (AP) occurrence while deconvolution at the most stringent threshold detects fewer false positive AP. Considering such features, we applied DeepSpike inference in the more dynamic-sensitive USN detection and information theory analyses. As synchronization analyses seek to identify highly synchronized cells and are prone to false positive bias, the deconvolution method was applied. We selected the highest threshold for deconvoluted spikes to maximumly reduced spike count and rule out false positives. In addition, we examined the performance of both methods by calculating false positive rates and accuracy values for all analyses. The false positive rate represents the proportion of negative signals that are incorrectly classified as positive. Accuracy represents the proportion of correctly classified spikes out of the total number of spikes. The low false positive rate (0.2 for DeepSpike and 0.1 for deconvolution) and high accuracy rate (0.8 for DeepSpike and 0.9 for deconvolution) validate the robustness of both methods.

With the spike-time information converted into binary series, each imaging frame yielded a multi-dimensional array consisting of 0 and 1. Based on inferred spike time series (binarized matrices consisting of 0 and 1, with 1 denoting one spike, and 0 denoting no spike, please see the diagram below), we counted the number of co-occurring spikes between a neuron (the target neuron) and any other neurons. For every imaging frame, if the target neuron was firing (“1”), we counted the spike number from all other neurons within that frame as the frame-wise synchronization count. If it didn’t fire (“0”), this frame is skipped. These frame-wise synchronization counts are added up across 2800 frames (90 s) to obtain synchronization counts for a single neuron.

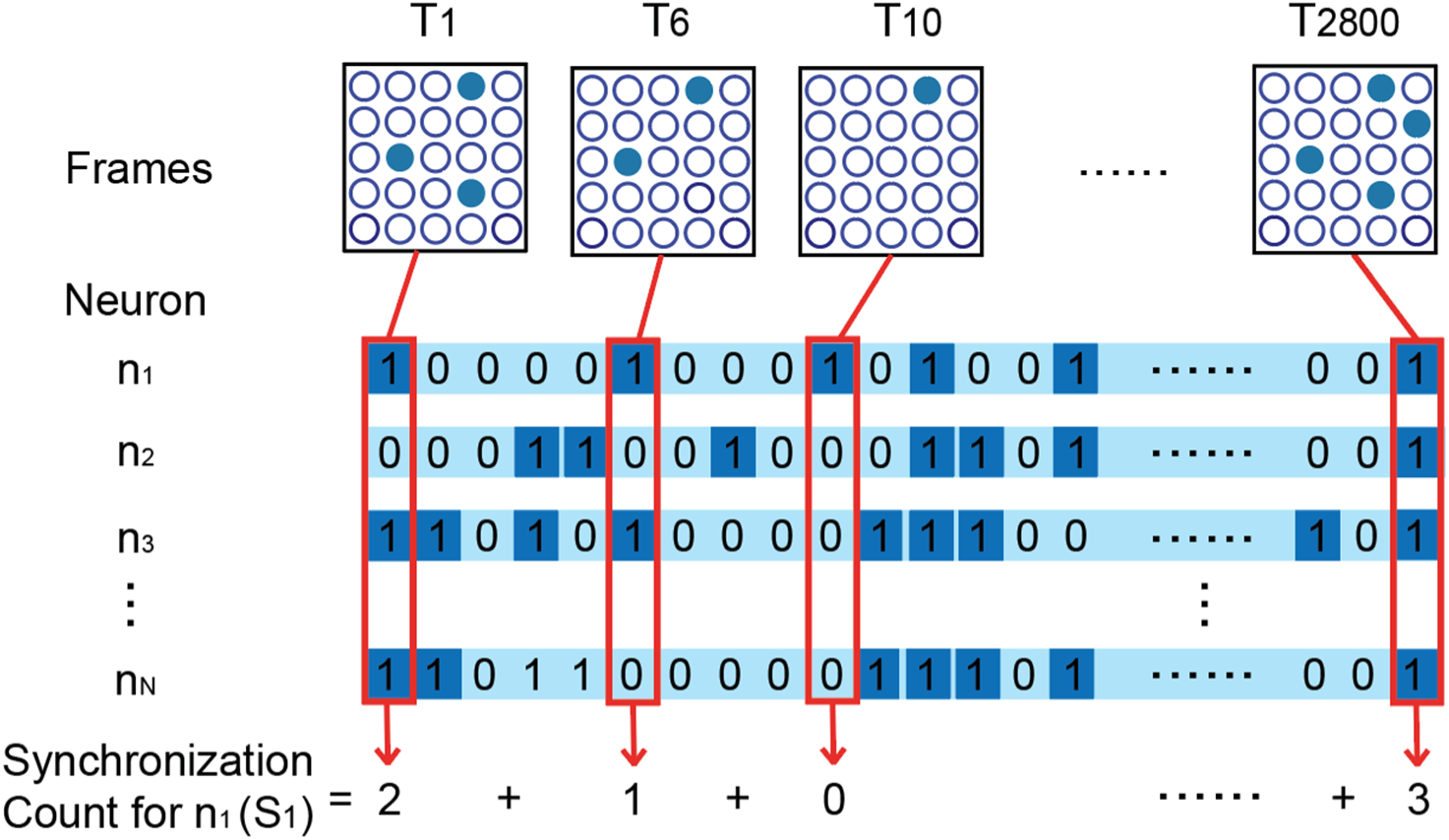

Next, the above process was repeated for all neurons in the imaging plane. Then the mean neuronal synchronization was taken as the total synchronization count S_mean_:

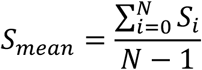

Where N = the number of all neurons in the field of view.

#### Synchronization normalization for chance firing interference

Since high firing rates could cause chance elevations in synchronization counts, we corrected the raw counts by subtracting randomly shuffled results (the shift predictor). These procedures were proceeded by shuffling the original spike-time series in order to randomly assign the spikes to different frames (please see the diagram below). The shuffling procedure was repeated for 1000 times to generate the final averaged shift predictor which was then subtracted.

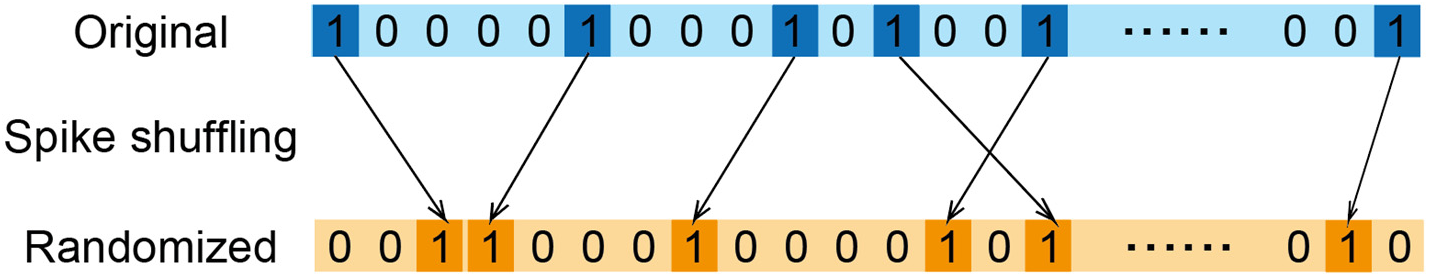

The corrected synchronization S_n_

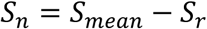

Where S_r_ is the average synchronization count calculated based on randomized ensemble firing patterns. Randomization process illustrated above was carried out 1000 times for each neuron, and all resulting S_r_ values was averaged to obtain the final S_r_ value in equation. S_n_ for each imaging timepoint was calculated following steps above, and the result was reported in Fig. 3c.

#### Validation of synchronization normalization method

Since high-firing cells may cause chance elevation in synchronization counts, we tested the validity of our normalization method using the following procedures. To test the effect of high-firing rate upon synchronization, we conducted a simulation experiment (Extended Data Fig. 10). Utilizing MATLAB program, we generated two artificial neurons and measured the impact of their firing rates upon synchronization. We artificially set different synchronization counts for these two neurons to serve as the “ground truth”. We then calculated normalized synchronization using between these two neurons using methods described above. We subtracted the “ground” truth from the calculated results to obtain error value of this method and assessed the effects of different sync. ratio on the error value (Extended Data Fig. 10b). The results showed that when the frequency of the neuronal activity (average firing rate) is less than 0.05, the original normalization method for synchronization impartially reflected the “ground truth” (Extended Data Fig. 10c). Since the average firing rate of actual neurons was close to 0.01 < 0.05, the original normalization method is applicable to our data.

To further corroborate that the synchronization analyses support our central claim, we also used another common normalization method of dividing the result by average neuronal firing rates. Specifically, for a neuronal ensemble that has N neurons, synchronization count of one neuron S at one imaging timepoints:

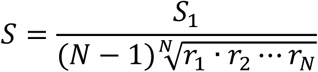

Where S_1_ denotes the raw synchronization count for neuron n_1_ (please see diagram Fig), r denotes the firing rate of a single neuron. The firing rate is calculated as follows:

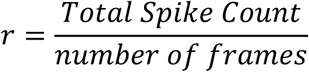

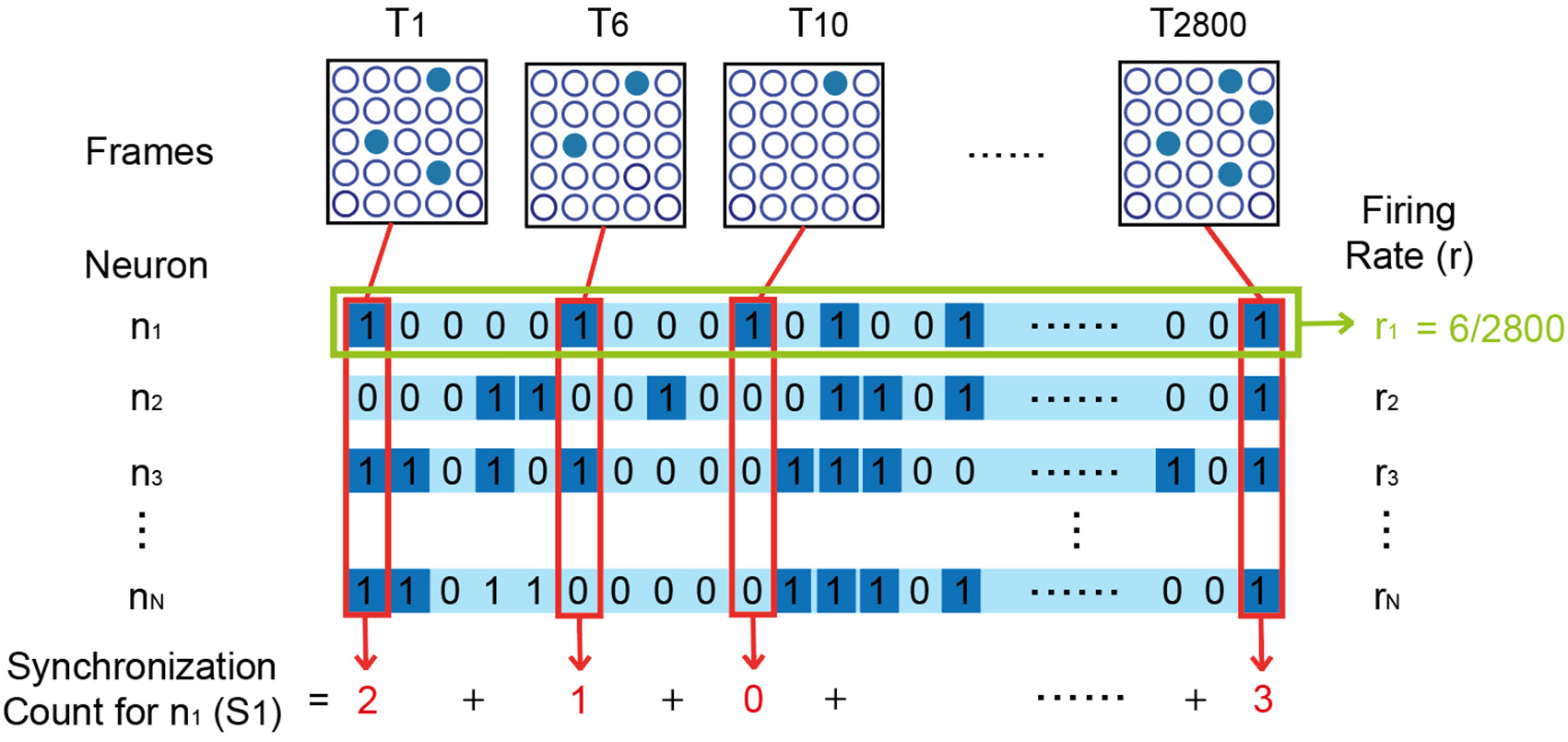

Then the sum of all neuronal synchronization was taken as the total synchronization count S_mean_:

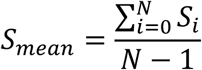

Where N = the number of all neurons in the field of view.

Lastly, corrected synchronization counts S_n_ was calculated as follows:

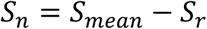

Where S_r_ is the average synchronization count calculated based on randomized ensemble firing patterns. Randomization process illustrated above was carried out 1000 times for each neuron, and all resulting S_r_ values was averaged to obtain the final S_r_ value in equation.

This calculation method of normalized synchronization counts still supported our key observation in Figure 3. That is, neurons that maintain high synchronization counts throughout the remote stage (d28-d63) fully included all USNs.

#### Definition of high sync. and low sync. neuron

Examination of synchronization counts for USNs revealed that they maintained high synchronization (sync. counts ≥ 3σ) for every remote imaging timepoint from d28 to d63 (Extended Data Fig. 7a). Based on this phenomenon, we defined high sync. neurons as neurons that maintained sync. counts ≥ 3σ for every remote imaging timepoint (28d, 39d, 45d, 49d, 52d, 59d, 63d). Neurons with sync. Counts < 3σ at any time point were defined as low sync. neurons.

We further tested this phenomenon in all GABAergic and non-GABAergic neurons using unsupervised clustering analysis. K-means clustering of standard deviation of Δ sync counts clearly showed that the high sync. neuronal group composed only of GABAergic neurons (Extended Data Fig. 7). Here Δ sync counts referred to the difference between the total sync. counts of each remote imaging timepoints (28d-63d) and the total sync. counts of before training baseline. Results from K-means clustering indicated that the existence of high and low sync groups is a phenomenon specific to GABAergic neurons (Extended Data Fig. 7).

#### Population vectors and ensemble similarity index calculation

Population vectors were calculated according to L. Carrillo-Reid et al.^85^ Neural activity patterns at a single imaging frame (33 ms time bin) can be written as an n-dimensional vector, with n denoting the total number of neurons in the imaging plane. For each frame, the activities of all neurons were described as a vector and each cell was seen as a variable of the vector. One vector thus represents the ensemble response at one moment in time. Cosine similarity calculates the difference in the spatial angle of the multi-dimensional vectors and helps illustrate changes in ensemble patterns over time or under different stimulation conditions. The calculation formula is as follows:

For two multi-dimensional vectors A and B

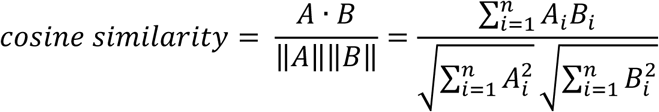

Where *A*_i_ and *B*_i_ denotes the i^th^ variable of vector A and B, respectively.

#### Complex network analysis

Weighted complex networks were used to describe the functional connections between the cells of the observed microcircuitry. Each cell was seen as a node of the complex network. We constructed the adjacency matrix of the complex networks from neural activity time series. Each element of the adjacency matrix (*W*) between a pair of cells denoted the count of how many times the two neurons fire simultaneously.

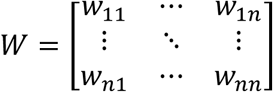

Where adjacency matrix (*W*) was symmetric. The diagonal element was set as zero. We adopted two criteria to describe the properties of the microcircuitry. Firstly, the values in the weighted adjacency matrix denotes the functional connections between cell pairs. A bigger *w_mn_* value indicates that the neuron pairs, *m* and *n*, are more tightly coupled in this observed microcircuitry. Secondly, the node strength (*NS_p_*) denotes the total count of simultaneous spikes between a target cell *p* and all other cells in the observed microcircuitry. *NS_p_* was calculated by

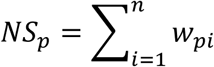

#### Topology analysis

Topology analysis was based on the complex functional network analysis and was used to describe the structural dynamics of different functional neuronal ensembles. To build the coactivity complex of the key neurons, we applied the “sliding coactivity integration window” approach. This analysis converts neural coactivity into a simplicial complex, whose spatial structure is determined by the relative timing of spikes in a neural population. Firstly, we identify the maximal simplexes that emerge within the first time period, based on the cell activity rates evaluated, and construct the corresponding input integration coactivity complex. Then we calculated the functional network for the subgroup of cells in different experimental statuses.

#### Joint information entropy and joint mutual information analyses

Joint information entropy and joint mutual information analysis were conducted using the Neuroscience Information Theory Toolbox. (https://github.com/nmtimme/Neuroscience-Information-Theory-Toolbox). For each neuron, spike results were converted to a binary activation time series composed of 1 and 0, with 0 denoting no spikes, and 1 denoting a spike. These discrete-time series (lasting 90 s) of DeepSpike yielded calcium spikes at different imaging time points were used to calculate joint entropy and mutual information. Joint entropy between each neuronal pair is calculated by the following formula:

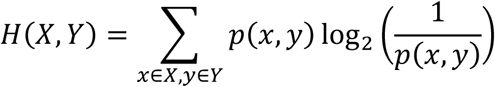

Where x denotes states (0 or 1, transcribed from DeepSpike inferred spike time series) of neuron X, and y denotes states of neuron Y. p (x = 1) denotes the calcium spike probability of neuron X, i.e., the ratio of the count of sampling points with calcium spike to the total sampling points. p (x = 0) = 1 – p (x = 1). p (x, y) denotes the joint probability distribution of neuron X and Y. p (x, y) has four states: p (x = 1, y = 1), p (x = 1, y = 0), p (x = 0, y = 1), p (x = 0, y = 0). The values of p (x, y) are determined by the spike time series of each neuron at each imaging time point. For example, p (x = 1, y = 1) denotes the ratio of the number of sampling points where both neurons X and Y have calcium spikes to the total number of sampling points.

Mutual information quantified the information that one neuron can provide about the second neuron and is calculated by:

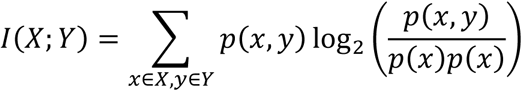

To test if the USNs demonstrates unique temporal dynamics in terms of information entropy, we compared their results to values generated from randomized controls. Randomized control samples were generated by shuffling the activation time series for each neuron. The spike order of each neuron was redistributed to randomly selected time points. The random experiments were repeated 10,000 times independently to get the average random measurements. To eliminate the influence of different neural activity strengths and improve the comparability of different experimental statuses, ratios between observed measurements and random measurements were calculated. To assess the training-induced changes in joint entropy and mutual information, we’ve also taken a second step of normalization, taking the ratio between measurements from each imaging timepoint and the measurements from before conditioning. Lastly, the population averages of normalized joint entropy and joint mutual information were calculated for various imaging time points.

#### Neural activity strength clustering analysis

Neural activity strength clustering analysis was used to find the key neuronal subgroup with high neural activities. Firstly, we calculated the spike count based on deconvolution and average amplitude of all calcium transients. Neural activity strength was associated with both the spike count and average amplitude. We set the baseline neural activity, denoting that the spike count and average amplitude were zero. We calculate each cell’s neural activity strength compared to the baseline using Mahalanobis distance. The Mahlanobis distance is defined as:

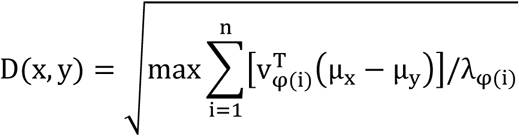

Whereas µ_x_ and µ_y_ are the average calcium signals from two neurons at each time session. λ_φ (i)_ is the eigenvalues of the covariance matrix of the union of x and y. V_φ (i)_ is the corresponding eigenvector and φ is a bijective mapping reordering indice of eigenvalues and eigenvectors. Second, we implemented hierarchical clustering for all cells based on their neural activity strength. The inner squared distance (minimum variance algorithm) was used when constructing an agglomerative hierarchical cluster tree.

#### Community Analyses

The multi-layer complex functional network analyses were performed by the software Gephi9.3 (open-source). The built-in community analyses detect similarities among connectivity data based on node and edge values in a network. In our case node information is composed of the average normalized calcium power for each neuron; edge information is composed of the normalized pairwise synchronization counts (see synchronization calculation) between each node. Multi-layer networks are constructed for neuronal ensemble activities under different stimulation settings at retrieval to visualize and assess their differences. Gephi assigns a modularity class index to each constructed network. The calculation equation is as follows according to the Gephi manual:

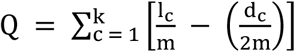

Where m is the total number of edges in the network; k is the number of communities; l_c_ is the number of connected edges within the community c; and d_c_ is the sum of the degrees of all the nodes in the community c.

#### Immunofluorescence

At the end of the 63-day experiment, histology of the imaged area was performed to assess neuronal health post long-term expression of rAAV2/9 viral tools. Mice were deeply anesthetized with tribromoethanol (0.02ml/g, intraperitoneally). Cardiac perfusion with 50 ml physiological saline and 25-50 ml 4% paraformaldehyde was performed sequentially. The brain was removed and immersed in 4% paraformaldehyde for 24 h at 4 °C. Then the brain was dehydrated in 40% sucrose for 3-6 days and frozen at -20 °C in optimal cutting temperature compound (OCT). Successive coronal sections of 30 μm were made with freezing microtome. The sections were mounted on slides with a mounting medium containing nucleic acid dye (DAPI, 4′,6-diamidino-2-phenylindole). Images were acquired using Nikon A1 confocal microscope with a 20 ×, 0.8 NA, and 40 ×, 0.6 NA objective lenses at 1024 × 1024 pixels. To count the number of GCaMP6f-positive cells and DAPI-positive nuclei, images of the ROI were acquired by ImageJ.

To examine the labeling efficiency and specificity of GABAergic neurons in GABA-Cre::Ai9 mice, we performed GAD67 immuno-staining. Coronal sections were prepared and mounted on slides following the above procedure. Then sections were first rinsed for 10 min × 3 times in PBS and blocked in PBS containing 10% donkey serum, and 3% Triton X at room temperature for 2 h. The primary antibody (#63080S, Cell Signaling Technology, In) was added and attached at 4 °C overnight. Then, sections were rinsed in PBS for 20 min and incubated with secondary antibodies (#A-31573, Thermo Fisher, Inc.) dissolved in PBS for 2 h at room temperature. Finally, sections were rinsed 3 times in PBS and mounted on slides with the nucleic acid staining mounting medium (DAPI, 4′,6-diamidino-2-phenylindole). For quantification, 7-10 z-stacks per animal were generated using a confocal microscope (Nikon, A1). Analysis by ImageJ revealed 100% signal co-localization with mCherry signals from report mice, consistent with the previous report.^41^

The neurotoxicity effect of long-term rAAV2/9 virus expression was assessed by TUNEL (terminal deoxynucleotidyl transferase-mediated dUTP nick-end labeling) Assay Kit (#64936, Cell Signaling Technology, Inc.). The toolkit enzymatically linked a fluorophore (640 nm) conjugated nucleotide to the 3’ end of fragmented DNA. Sections were rinsed two times in PBS for 5 min each, permeabilized in PBS-TB for 30 min, and then washed again two times in PBS for 5 min each. Then sections were incubated in 100 µL TUNEL Equilibration Buffer for 5 min. 50 µL of TUNEL reaction mix (1 µL of TdT Enzyme added to 50 µL of TUNEL Reaction Buffer) was added to each slide. Then sections were incubated for 60 min at 37°C protected from light. Post incubation, sections were rinsed three times in PBS-TB for 5 min each. The slides were finally covered with a mounting medium containing nucleic acid dye (DAPI, 4′,6-diamidino-2-phenylindole). 7-12 z-stacks per animal were acquired using a confocal microscope (Nikon, A1). Co-localization of GCaMP6f, C1V1(T/T), eNpHR3.0 and TUNEL fluorescence was examined in ImageJ (Extended Data Fig. 2). Analysis revealed a 1.29% apoptosis rate in GCaMP6f+ cells. Among all GCaMP6f+ cells, 11.93% of TUNEL+ cells were found at the injection injury site (30-50 μm radius), 1.08% near injury (50-130 μm radius), and 0.07% away from injury. To test whether apoptosis was caused by physical injury of the glass syringe, we inserted needle without the virus and counted TUNEL+ neurons (Extended Data Fig. 2c). We found that the majority of TUNEL+ cells resided at the injury site, suggesting that physical injury was the main cause of apoptosis. No significant neurotoxicity effect was found for long-term rAAV2/9 viral tools expression.

#### In vivo whole-cell electrophysiology

In vivo whole-cell electrophysiology was performed under two-photon imaging. Borosilicate glass pipettes (6–8 MΩ) were pulled by horizontal micropipettes puller (P-97, Sutter Instrument, USA) and filled with an intracellular solution (135 mM potassium methylsulfate, 8 mM NaCl, 10 mM HEPES, 2 mM Mg-ATP, 7 mM phosphocreatine, 0.01 mM d-Bicuculline, 0.02 mM Alexa Fluor 594, with 7.3 pH, 316 mOsm). Pipettes were positioned using a micromanipulator (MP-225, Sutter Instrument, USA). Data was recorded with a Axopatch-700B amplifier (Molecular Devices, Palo Alto, CA) and was low-pass filtered at 2 kHz and acquired at 5-10 kHz. Collected data was processed with the pClamp10.1 software and analyzed using Clampfit 10.1 (Molecular Devices, Palo Alto, CA).

### Statistical analysis

All statistical analyses were performed using GraphPad Prism software (GraphPad Software, Inc., La Jolla, California, USA). The data were presented as the mean ± SEM (Standard error of mean). Parametric analyses were performed using one-way ANOVA with Tukey post-hoc tests. Nonparametric analyses were performed using Friedman test with Dunn’s correction for multiple comparisons. Statistical significance threshold was set at 0.05, and significance levels were presented as \**P* < 0.05, \*\**P* < 0.01 in all figures unless otherwise noted.

## Extended Data

**Extended Data Fig. 1.**
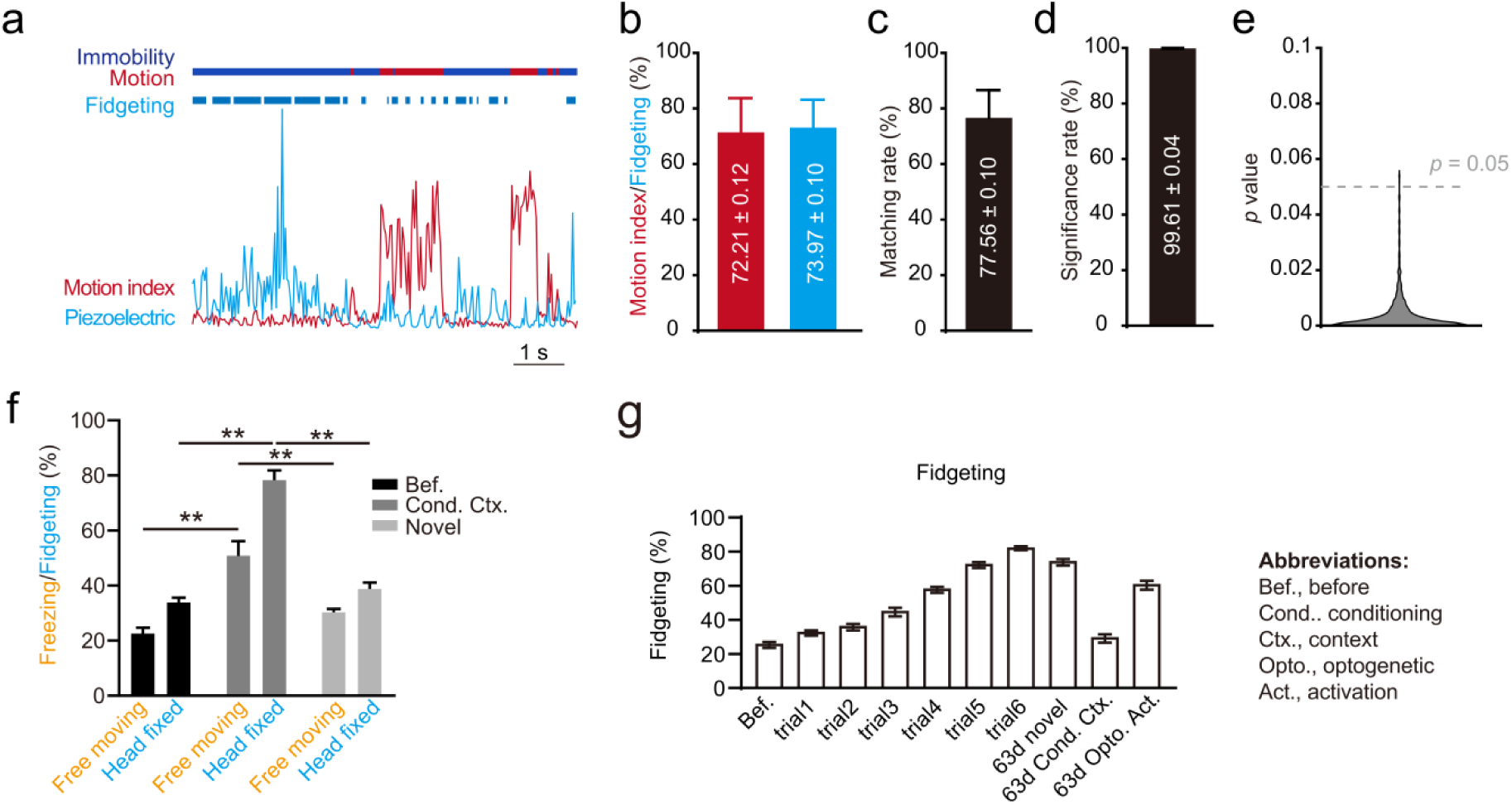
Behavioral specificity of piezoelectric signals. **a,** Validation of conditioned behavioral response assessed by video analyzation and piezoelectric signals respectively. Note, this figure was borrowed from Fig. 1c for clearer presentation of other data in this supplementary figure. **b,** Comparable percentage of fidgeting and immobility time from sample shown in a. **c,** Matching rate between labeled fidgeting and immobility from the sample in A. **d,** Significance rate (P < 0.05) of 1000 t-tests based on simulated fidgeting score of remote memory retrieval and before conditioning. 99.61% significance rate indicated that 99.61% of p-values were smaller than 0.05 in all simulations. **e,** Remote memory retrieval induced significantly higher fidgeting compared to before even for simulated p-values (overall distribution < 0.05). **f,** Side by side comparison of fear conditioning results measured by percentage of freezing (free moving) and fidgeting (head fixed) respectively. Free moving CFC: before = 22.50 % ± 2.22 %, old = 50.75 % ± 5.39 %, novel = 30.25 % ± 1.32 %, old vs. before, P = 0.0007 < 0.001, novel vs. before, P = 0.299 > 0.05, old vs. novel, P = 0.006 < 0.01; Head fixed CFC fidgeting%: before = 33.75 % ± 1.89 %, old = 78.25 % ± 3.64 %, novel = 38.75 % ± 2.39 %, old vs. before, P < 0.0001, novel vs. before, P = 0.435 > 0.05, old vs. novel, P < 0.0001, n=4 animals. **g,** Conditioning trial-averaged fidgeting response demonstrated successful formation of associative memory. Before: 25.33 ± 1.76, trial 1: 32.25 ± 1.53, trial 2: 35.75 ± 1.88, trial 3: 44.58 ± 2.57, trial 4: 57.58 ± 1.77, trial 5: 72.08 ± 1.68, trial 6: 82.00 ± 1.16, 63d trig: 73.67 ± 1.90, 63d novel: 29.08 ± 2.52, 63d opto: 60.25 ± 2.61; trial 1 v.s. Before: *P = 0.30 > 0.05*, trial 2 v.s. Before: *P = 0.0121 < 0.05*, trial 3 v.s. Before: *P < 0.01*, trial 4 v.s. Before: *P < 0.01*, trial 5 v.s. Before: *P < 0.01*, trial 6 v.s. Before: *P < 0.01*, 63d trig. v.s. Before: *P < 0.01*, n = 12 animals, one-way ANOVA, mean ± s.e.m.

**Extended Data Fig. 2.**
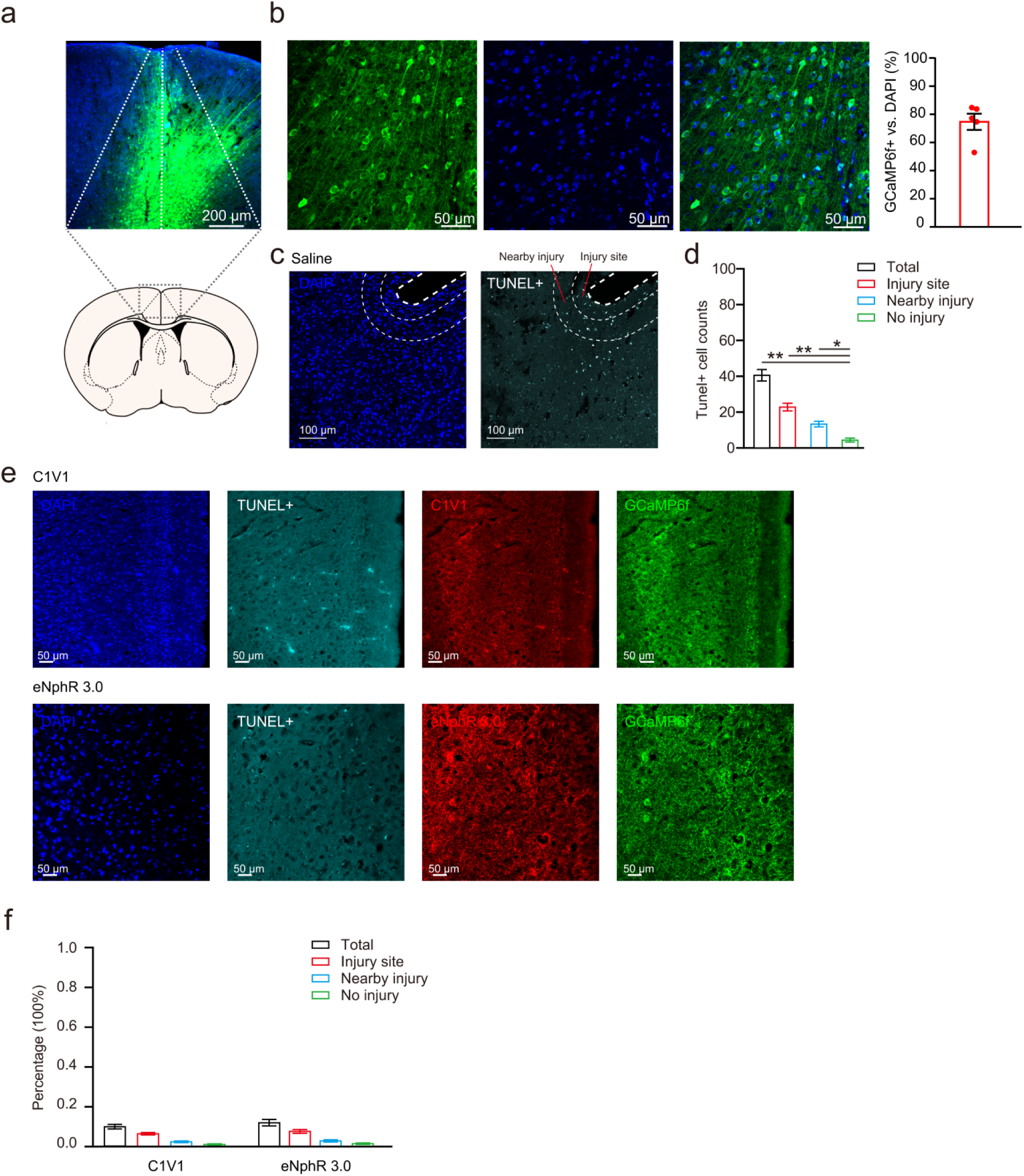
Verification of injection, viral expression and neuronal health post long-term expression of rAAV2/9. **a,** Location of rAAV2/9-hsyn-GCaMP6f injection and expression in right anterior cingulate cortex (rACC, AP: 0.26 mm, ML: -0.35 mm, DV: 0.30 mm). **b,** Histology of GCaMP6f positive cells (green) and all ACC neurons (blue, DAPI stained). Numbers of cells were manually counted and percentage of GCaMP6f + cells versus DAPI stained cells was 73.51 ± 5.67 (Mean ± SE). Scale bar: 50 μm. **c,** Apoptosis near injection location assessed by TUNEL assay (n=7 slices from 2 animals). The injury site was defined by 30-50 μm radius around the needle lesions; “nearby injury” was defined by 50-130 μm radius. **d,** Total number of TUNEL+ cells in 12 slices from 3 animals. Total = 40.58 ± 3.18 number of cells, injury site = 22.83 ± 2.14, nearby injury = 13.33 ± 1.53, no injury = 4.42 ± 1.16. Total vs. No injury, P < 0.0001, injury site vs. No injury, P < 0.0001, nearby injury vs. No injury, P =0.03 < 0.05, one-way ANOVA. **e,** Location of rAAV2/9-hsyn-C1V1, rAAV2/9-hsyn-eNpHR3.0 and GCaMP6f injection, expression and TUNEL assay signals in right anterior cingulate cortex. **f,** Percentage of number of TUNEL+ cells in 12 slices from 3 animals of each group. The injury site was defined by 30-50 μm radius around the needle lesions; “nearby injury” was defined by 50-130 μm radius; “no injury” was defined by the rest of the imaged areas. C1V1: Total = 10.01 % ± 1.16 %, injury site = 6.50 % ± 0.62 %, nearby injury = 2.46 % ± 0.42 %, no injury = 1.06 % ± 0.38 %. eNphR3.0: Total = 12.10 % ± 1.62 %, injury site = 7.66 % ± 0.93 %, nearby injury = 2.92 % ± 0.53 %, no injury = 1.53 % ± 0.38 %.

**Extended Data Fig. 3.**
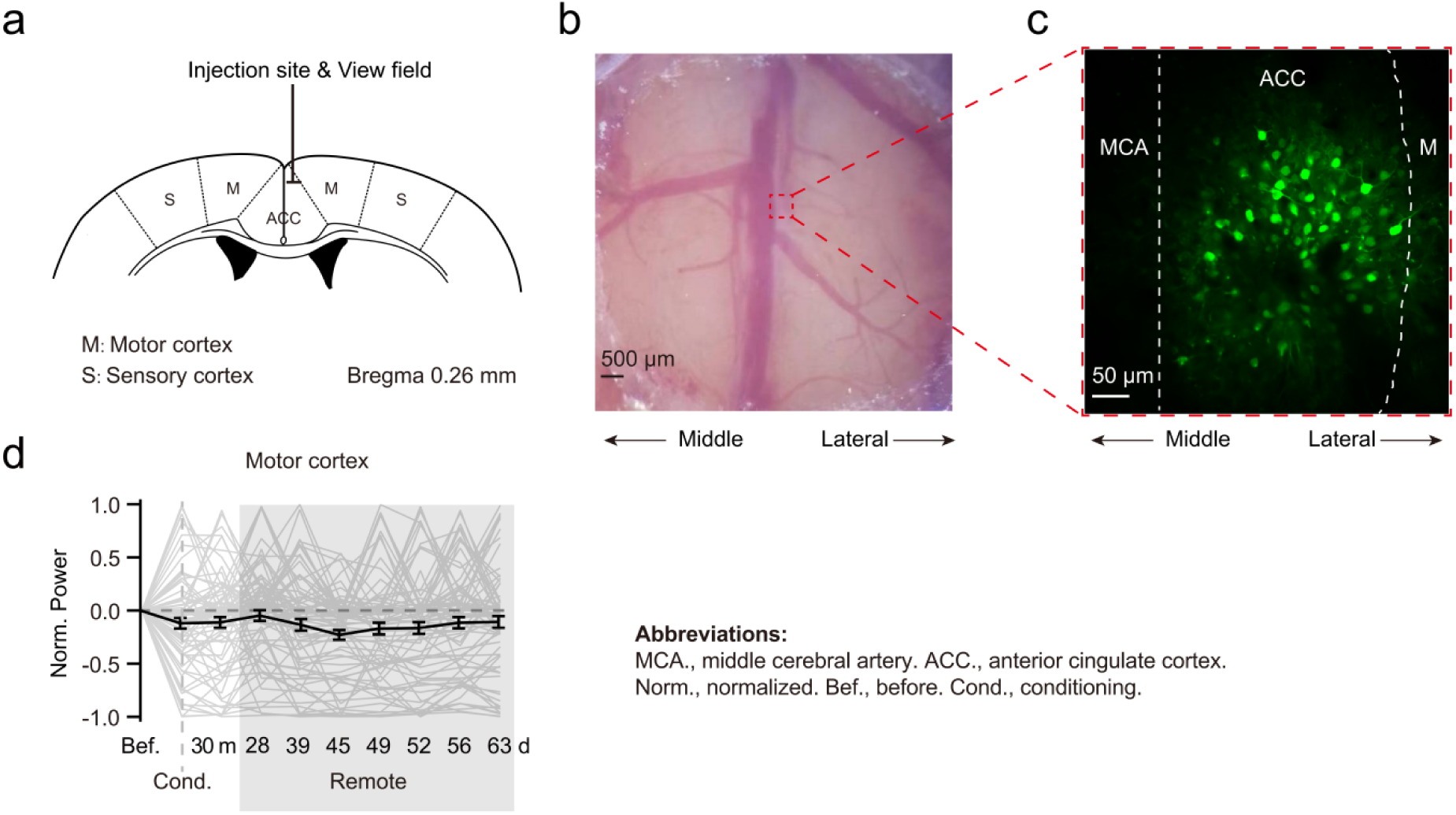
Corroboration of ACC targeting and region-specificity of USNs. **a,** FOV location on the mouse brain atlas, coronal section at 0.26 mm anterior to the bregma. **b,** Photo of the cranial window. Location of the FOV marked by red dotted lines. **c,** Sample FOV (averaged across 2800 frames). Location of the middle cerebral artery (MCA), ACC, and the motor cortex (M) were marked by white dotted lines. Note, this figure was borrowed from Fig. 1e for clearer presentation of other data in this supplementary figure. **d,** Absence of USNs in the neighboring motor cortex. No neurons in neighboring motor cortex maintained persistent above-zero average calcium power at remote time points (28-63d), indicating that USNs could not be identified in the motor cortex. Black lines, average calcium power. Gray lines, individual neuronal calcium power. Remote stage results were shaded in gray.

**Extended Data Fig. 4.**
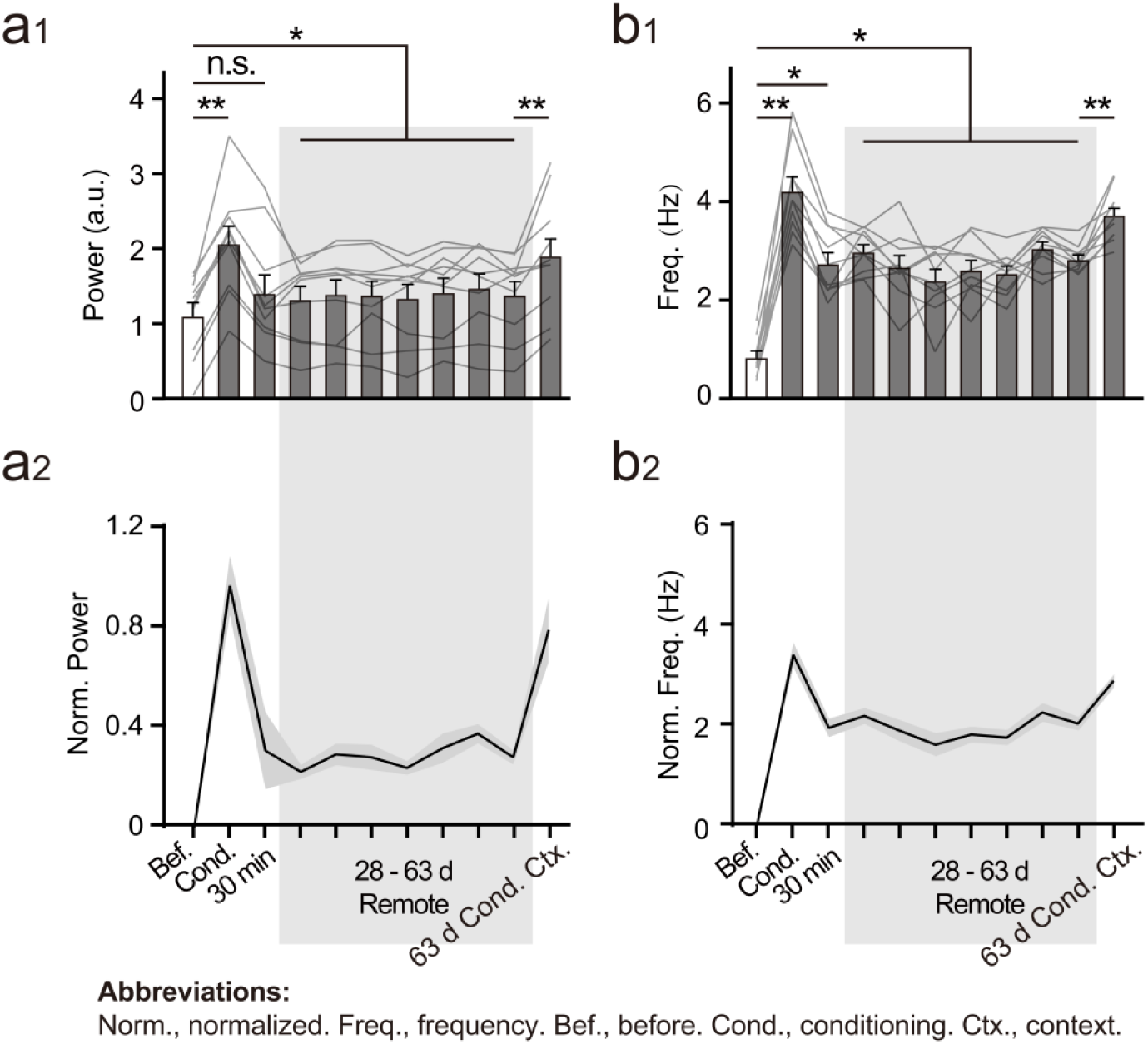
USNs remain in an intermediate activity level until being further activated upon retrieval. Comparing retrieval and default activities revealed the convergent dynamic feature of these neurons across subjects. Activities of USNs during retrieval (shown in red box) were much higher than default potentiated activities during the remote stage, indicating that these cells remain in an intermediate state until being further activated by retrieval cues. a1&b1, original values of calcium power and spike frequency. a2&b2, normalized calcium power and spike frequency showing the difference between post-training and before training activities.

**Extended Data Fig. 5.**
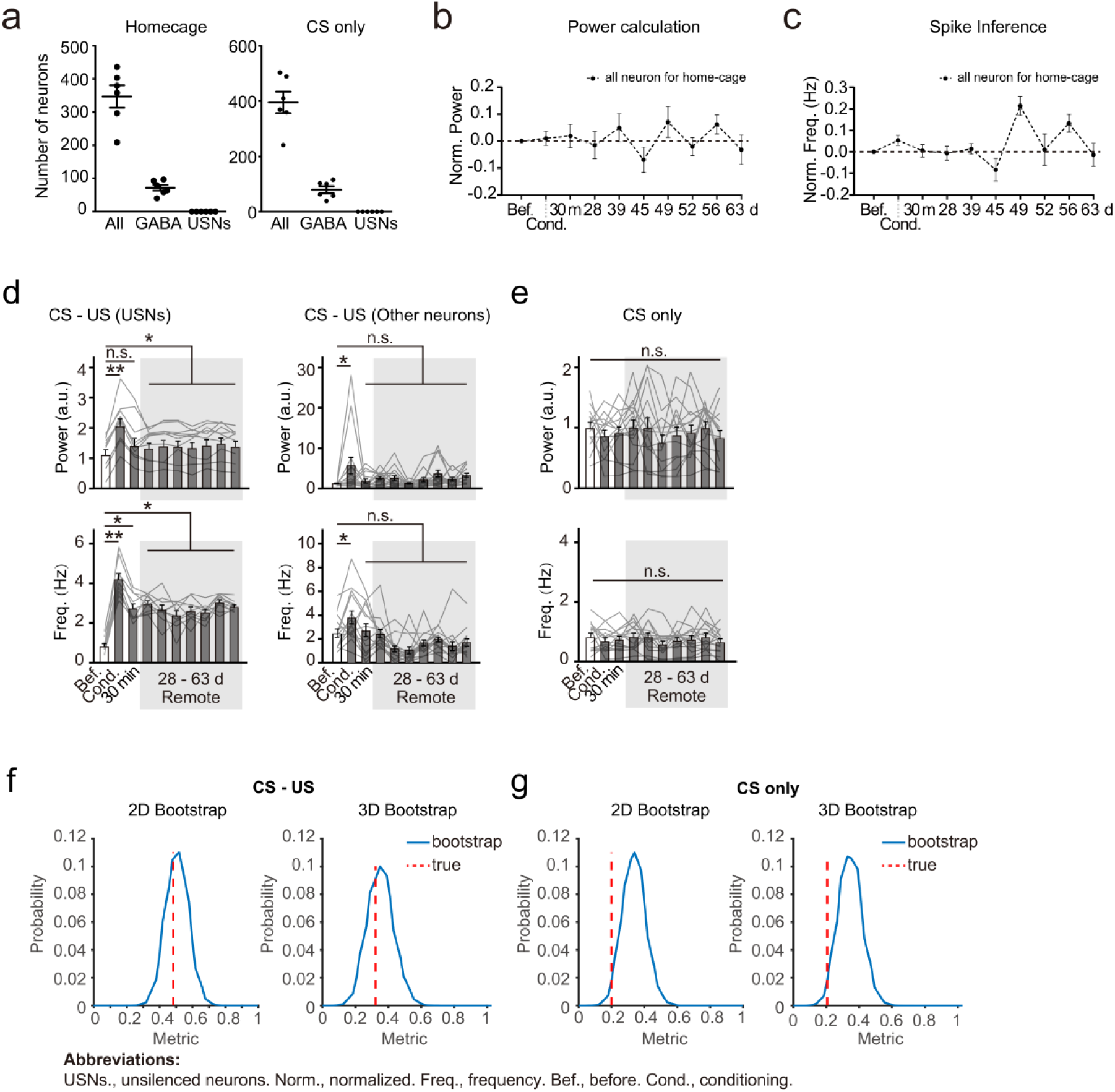
Absence of USNs in CS only group. **a,** Distribution of neuronal types in control groups. No USNs identified for CS-only and home caged groups. Home-caging: All neurons: 346.83 ± 33.59, GABA neurons: 72.00 ± 8.89, USNs: 0.00 ± 0.00, CS only: All neurons: 395.17 ± 39.33, GABA neurons: 80.54 ± 12.54, USNs: 0.00 ± 0.00, mean ± s.e.m. **b,** Normalized calcium power dynamics in all neurons in home caged groups. Stochastic changes around before-conditioning baseline were observed at remote stage. **c,** Normalized calcium spike counts in all neurons in home caged groups confirmed results of power calculation. Spikes were inferred using DeepSpike method. **d,** USNs in CS-US group showed more elevated average power and spike frequency during remote stage compared to other neurons. Gray lines, results from individual neurons. Gray area, remote memory stage. USNs, before vs. cond. P < 0.001; before vs. remote, P < 0.05; L1, before vs. cond. P < 0.01, before vs. 30 min, P < 0.05, before vs. remote, P < 0.01; Other neurons, before vs. cond. P < 0.05, each timepoint’s value was compared to before’s value using nonparametric Friedman test with Dunn’s correction for multiple comparisons, n = 52 neurons for USNs and other randomly selected neurons respectively from 12 animals). **e,** No distinct pattern in average power and spike frequency can be identified for CS-only neurons. Power: P = 0.999 > 0.05, spike frequency: P = 0.999 > 0.05, n=78 neurons in 5 animals, one-way ANOVA, Dunn’s multiple comparisons test. **f,** Bootstrap analysis for 2D and 3D imaging in CS-US group. Overlap of estimator (red) and distribution (blue) in CS-US group indicate the presence of USNs. n = 5 animals. Blue lines, the bootstrapped curve of neuronal spikes at all time points. Red dotted line, the average probability value of finding an “USN” in 1000 randomly shuffled imaging results. **g,** Bootstrap analysis for 2D and 3D imaging in CS-only group. Separation of estimator (red) and distribution (blue) in CS-only group suggest that USNs were absent.

**Extended Data Fig. 6.**
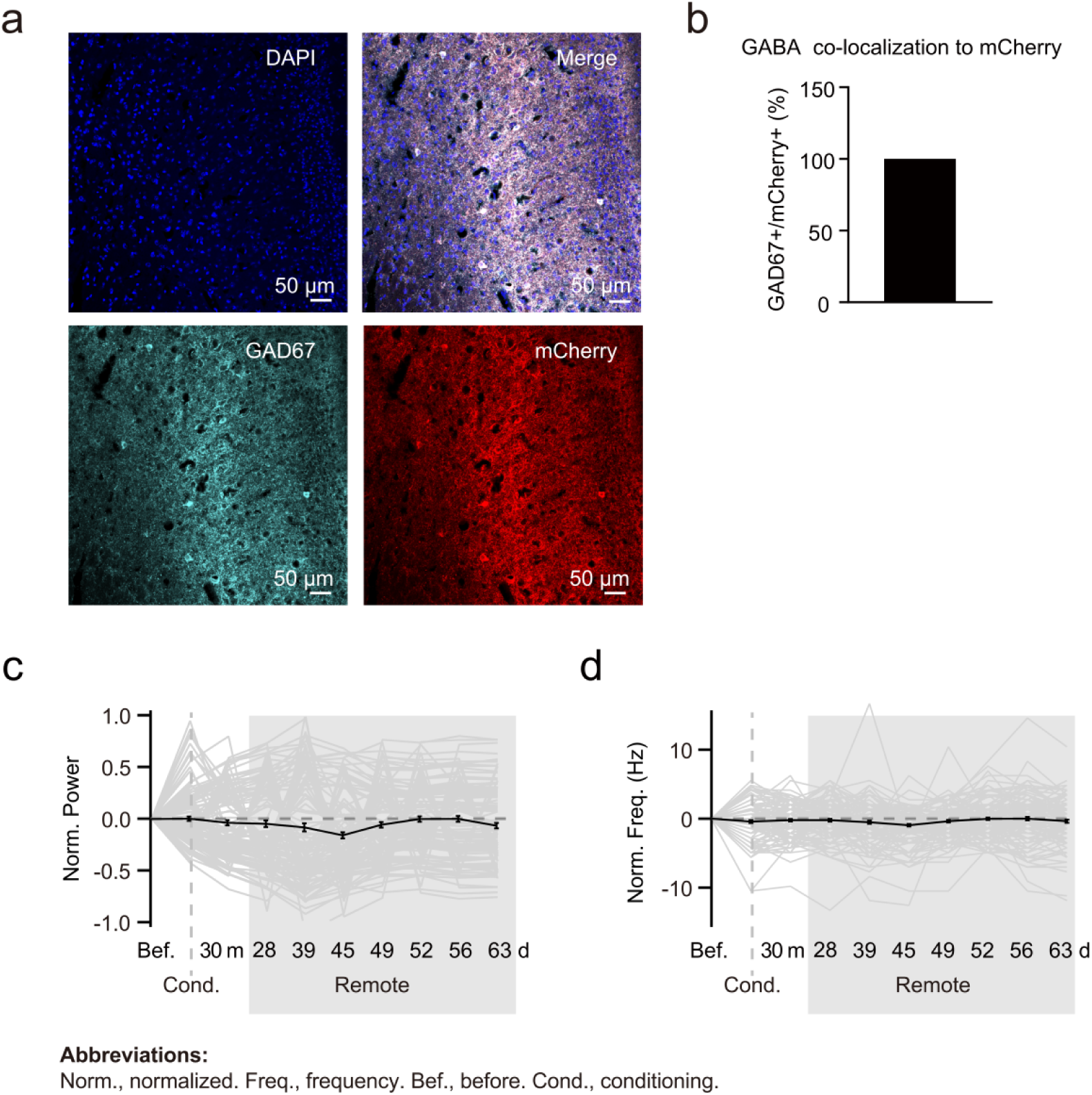
GABAergic labeling efficiency in GABA::Ai9 mice. **a,** Histology evaluating report specificity and accuracy of GABA::Ai9 mice reporters. **b,** GAD67+ cells and mCherry signals show 100% co-localization in the ACC region. GAD67+ cells / mCherry+ cells = 100%, n = 6 ROIs from 6 slices of 2 animals. **c,** Activities of all GABAergic neurons fluctuate around baseline during remote memory stage, as opposed to USNs. Values at each imaging timepoints were normalized by subtracting the before conditioning baseline value. Black lines, average normalized power. Gray lines, individual neuron normalized power. Gray area, remote memory stage. n = 113 neurons from 1 sample animal. **d,** Normalized spike frequency of all GABAergic neurons.

**Extended Data Fig. 7.**
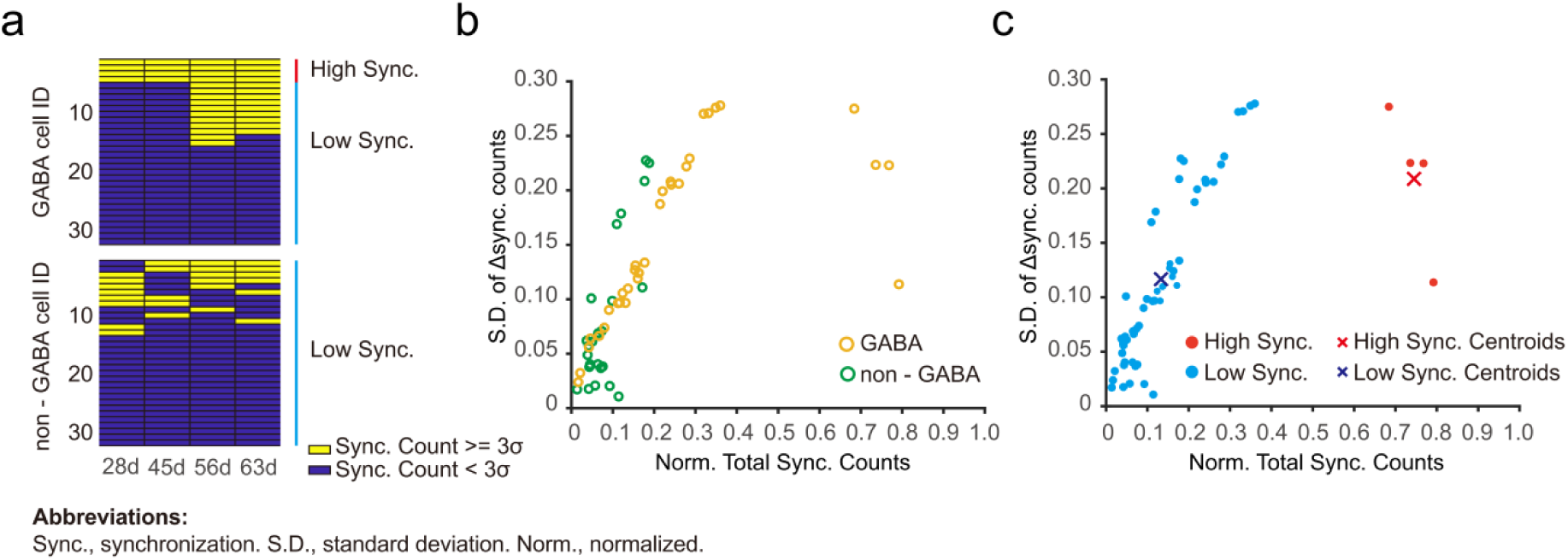
Two neuronal groups with distinct synchronization stability were identified in GABAergic neurons, but not in non-GABAergic neurons. **a,** No high sync. neurons (red bar) could be identified in non-GABAergic cell groups. Yellow indicated that the neuronal sync. count ≥ 3σ at a specific timepoint. For a cell to be identified as “high sync.”, all four remote timepoints (28d-63d) needs to be yellow. Blue indicated low sync. neurons (sync. count < 3σ). **b,** Distinction between GABAergic and non-GABAergic neurons in distribution of Δ sync. counts’ standard deviation and normalized total sync. counts. **c,** Unique cluster of high sync. neurons existed only in GABAergic neurons, but not non-GABAergic neurons. K-means values were calculated by the Euclidean distance.

**Extended Data Fig. 8.**
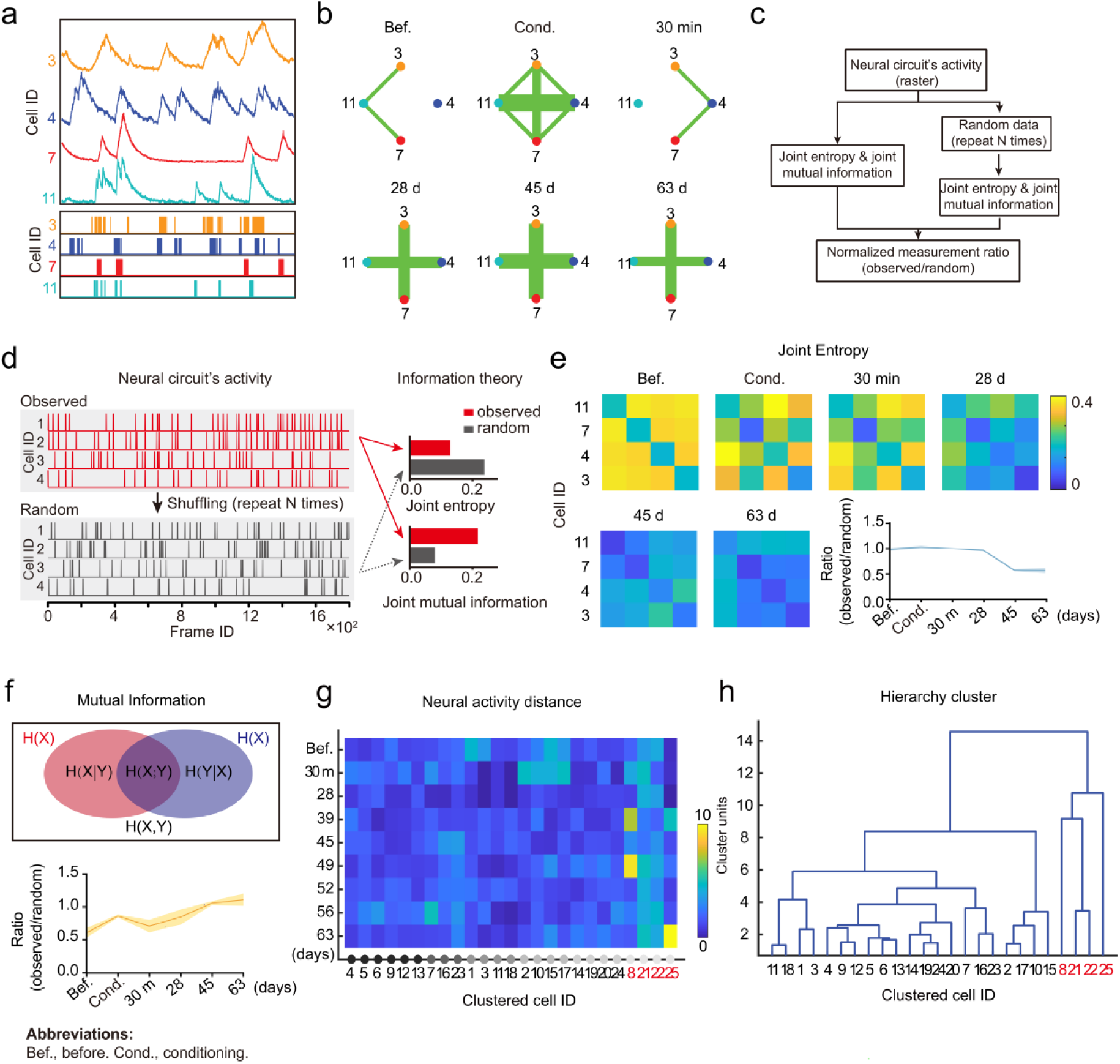
USNs constitute a stable topological network for efficient information processing. **a,** Calcium spikes inferred from raw calcium transients using DeepSpike. Colored numbers indicate USN IDs. **b,** Topological map of USN ensembles at different stages of contextual memory formation. Topology reflects how synchronized firings are processed by the readout neurons. Node colors indicate cell IDs in panel a. Line thickness indicates the stabilization probability of the functional connectivity between each node (for calculation methods, see supplementary materials). **c,** Flow chart of joint information entropy and mutual information analyses. **d,** Observed unsilenced ensemble activities show higher capacity for information processing compared to that of the randomized control. Left, a randomized control was generated by shuffling neuronal spike time series N times. Right, diagram of parameters calculated from real data (observed) and randomly generated samples (random). **e,** Decreased joint information entropy in USNs supports efficient information processing at remote stages. Top and bottom left, joint information entropy of USNs (n = 4 neurons, 1 animal). Heatmaps mirrored along the diagonal line. Bottom right, temporal changes show decreases in averaged normalized joint entropy at the remote stage (Before: 0.987 ± 0.021, Conditioning: 1.032 ± 0.024, 30 min: 1.002 ± 0.001, 28 d: 0.970 ± 0.018, 45 d: 0.576 ± 0.020, 63 d: 0.571 ± 0.053, n = 5 animals, mean ± s.e.m.). Low information entropy values at remote time points indicate high nonlinear correlations in pairwise neuronal activities. Normalization is performed by taking the ratio between the observed sample results and random sample results. **f,** Increased joint mutual information of USNs at the remote stage supports efficient information processing. Top, diagram showing the calculation method for mutual information. Bottom, temporal changes in average normalized joint mutual information (Before: 0.611 ± 0.074, Conditioning: 0.867 ± 0.022, 30 min: 0.714 ± 0.094, 28 d: 0.852 ± 0.114, 45 d: 1.064 ± 0.029, 63 d: 1.114 ± 0.094, n = 5 animals, mean ± s.e.m.). **g,** Mahalanobis distance between ensemble activities at each imaging time point and activities before training. The results are from all observed GABAergic neurons. The color bar indicates the distance value. Cells that possess large distances underwent drastic activity shifts during remote stages compared to before conditioning (red, USNs: 8, 21, 22, 25). n = 24 neurons in 1 animal. **h,** Hierarchical clustering analysis indicates that USNs (red, 8, 21, 22, 25) are clustered and display similar activity patterns. Clustering was based on Mahalanobis distances further revealed the shortest within-group distance among USNs, indicating their convergence in temporal activity dynamics.

**Extended Data Fig. 9.**
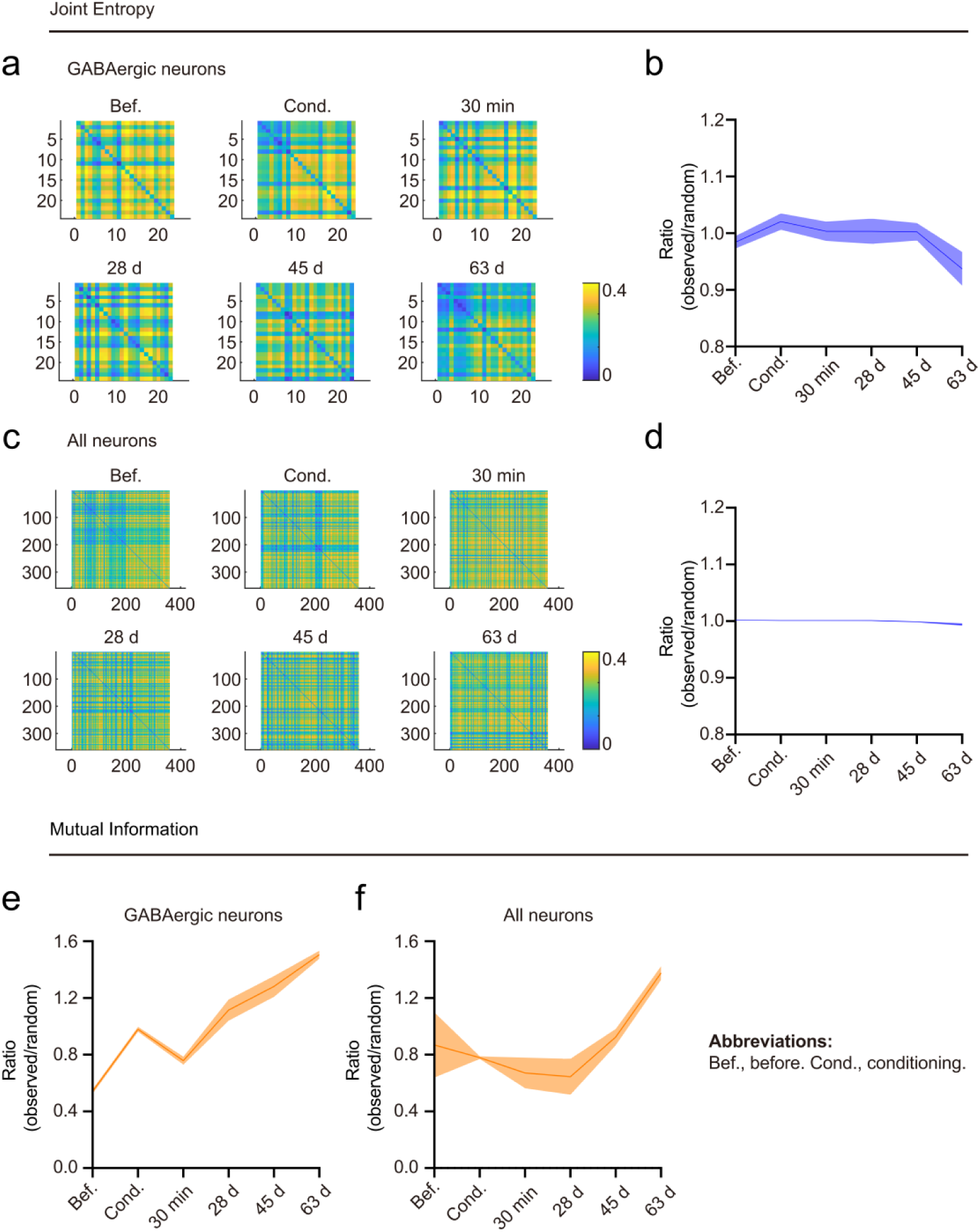
Joint information entropy and mutual information dynamics of all GABAergic neurons and the entire imaged population. **a,** Joint information entropy for all GABAergic neurons imaged across different time points in 1 animal. Vertical axis denotes cellular ID number. **b,** Temporal changes in average normalized joint information entropy for all GABAergic neurons. Mean ± s.e.m. Normalization was performed by taking ratios between observed and randomly shuffled control samples’ results. Similar to results for USN ensembles, normalized joint information entropy showed a decreasing trend at the remote stage. **c,** Joint entropy for all neurons (GABAergic and non-GABAergic neurons) imaged across different time points in 1 animal. **d,** Temporal changes in normalized joint information entropy for all neurons. The small variance in entropy derived for collective neuronal activities indicate strong stochasticity in their activation patterns. **e,** Temporal changes in normalized mutual information for GABAergic neurons. Mean ± s.e.m. **f,** Temporal changes in normalized mutual information for all neurons. Mean ± s.e.m.

**Extended Data Fig. 10.**
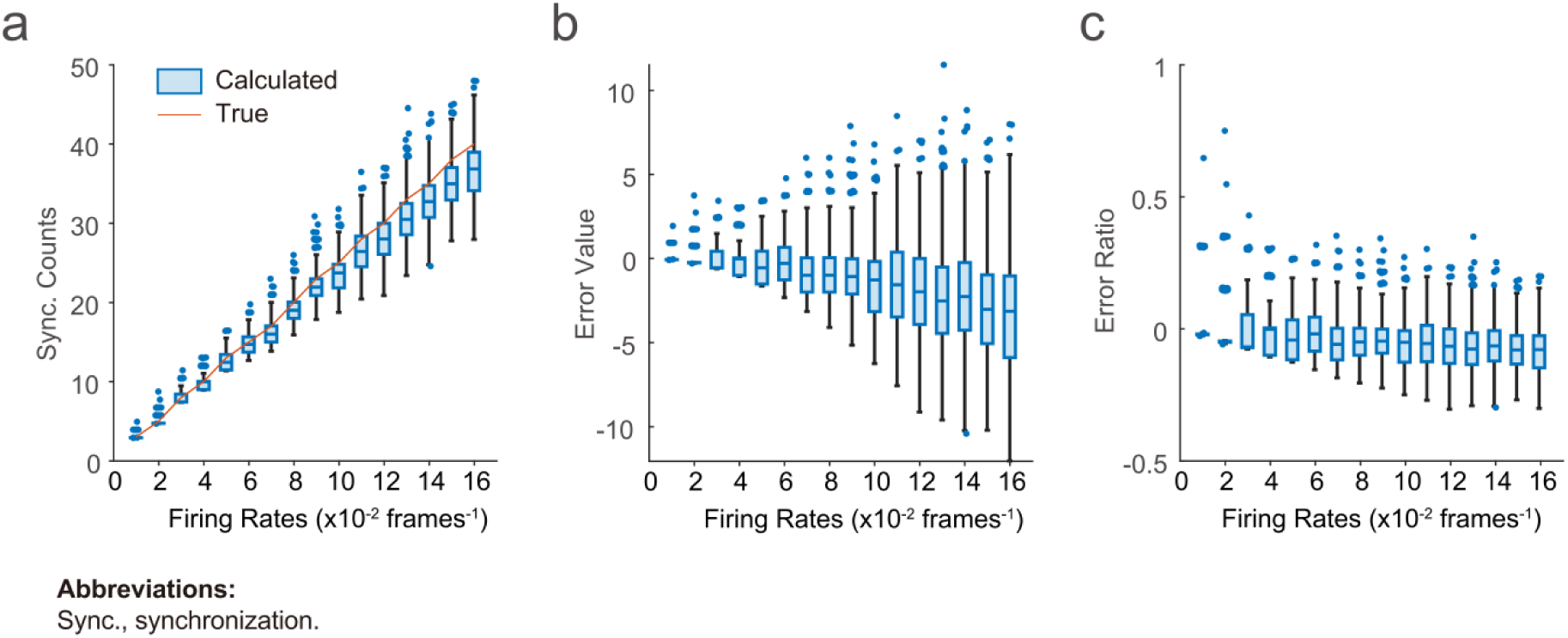
Assessment of synchronization normalization method by simulation experiment. **a,** Sync. counts value calculated and normalized with original methods was comparable to ground truth sync. counts at all firing rate levels. Firing rate was obtained from firing counts of two artificial neurons within one imaging frame. **b,** Smallest error value was obtained at firing rates smaller than 0.01. Error value was calculated by taking the difference between calculated sync. counts and ground truth sync. counts. **c,** The smaller the firing rate, the smaller the error ratio in the 1000 simulations. For firing rate smaller than a certain level (e.g. 0.01), error ratio in synchronization estimation was neglectable. Error ratio was calculated by dividing the error value with calculated sync. counts.

**Extended Data Fig. 11.**
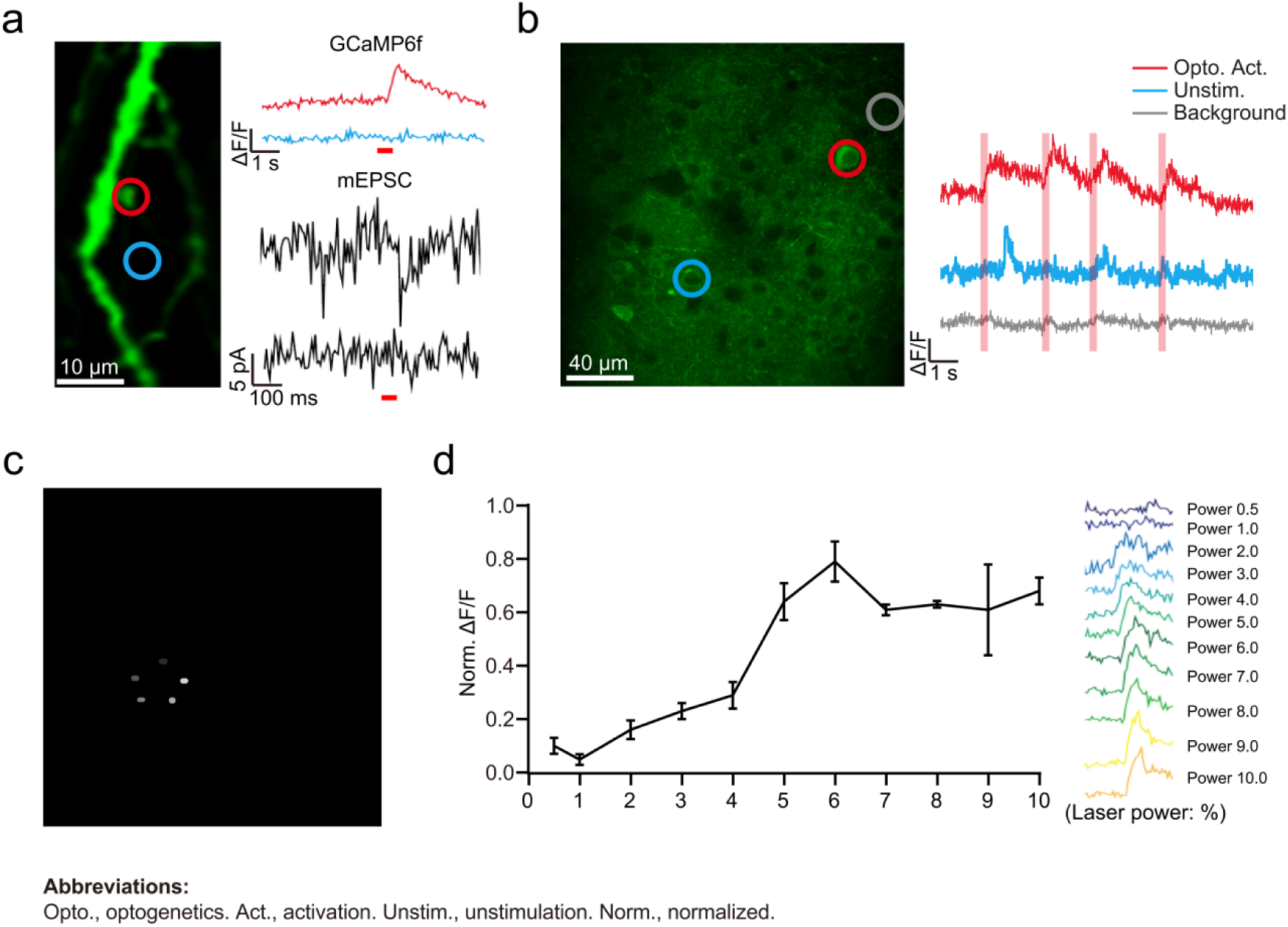
Precision verification of two-photon holographic optogenetic stimulation. **a,** Holographic stimulation by galvano-galvano lens of a single spine (red) created mEPSPs while a proximal stimulation (blue) did not. Left, selected stimulation ROIs. Top right, calcium transients (ΔF/F) for the bouton (red) and control position (blue) respectively post stimulation. Bottom right, mEPSP recorded at the soma via *in vivo* patch-clamp. Red bar indicates light stimulation. **b,** Effect of stimulation in an example imaging field. Left, stimulation ROIs where stimulated neuron was circled in red, unstimulated neuron in blue and control ROI containing no neurons circled in grey. Right, GCaMP6f fluorescence transient (ΔF/F) trace for respective ROIs in b. Bar indicate stimulations. **c,** Simultaneous stimulation of 5 ROIs on a fluorescence plate using the SLM method. **d,** Line chart of averaged calcium signal showing increase for stimulation at different power levels. Right, ΔF/F trace for each power level.

**Extended Data Fig. 12.**
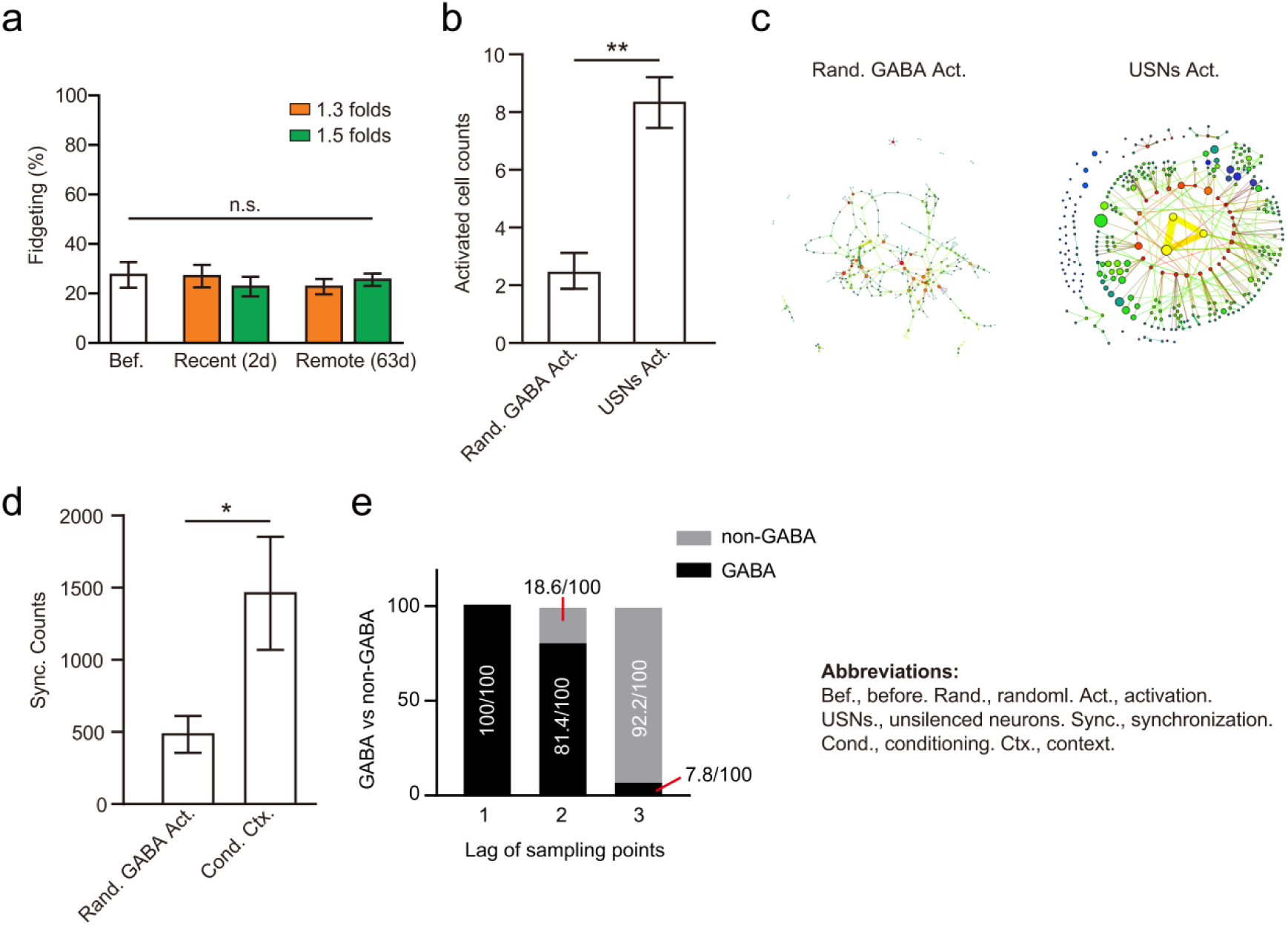
Random GABAergic cell stimulation failed to retrieve remote memory and memory-related hierarchical ensembles. **a,** Random activation of memory-related GABAergic cells failed to retrieve remote memory. Memory-related GABAergic cells were defined by neurons with increased activities during encoding compared to before training baseline. 1.3 folds and 1.5 folds referred to neurons with respective 30% and 50% activity increment during conditioning stage compared to before training baseline. 1.3 folds: before = 27.25 ± 5.17, recent = 27.00 ± 4.55, remote = 22.75 ± 3.97, recent vs. before, P = 0.94 > 0.05, remote vs. before, P = 0.74 > 0.05. 1.5 folds: recent = 22.75 ± 3.07, remote = 25.50 ± 2.50, recent vs. before, P = 0.99> 0.05; remote vs. before, P = 0.96 > 0.05, n=4 animals, one-way ANOVA. **b,** Fewer neurons fired in the first frame (33ms) after holographic stimulation of random GABAergic neurons compared to USNs. USNs Act. = 8.33 ± 0.88, random GABA Act. = 2.50 ± 0.62, P = 0.0003 < 0.001, n = 6 animals, two-tailed t test. **c,** Sparser evoked downstream pattern by random GABAergic neurons stimulation compared to USNs. **d,** Weaker functional connection between downstream neurons under random GABAergic stimulation compared to natural context recall. Random GABA stim. = 482.72 ± 127.37, natural context = 1461.02 ± 391.16, P = 0.0387 < 0.05, n = 6 animals, two-tailed t-test. Functional connection was measured by count of synchronization events (sync. counts) between neurons within the first frame (33ms) after opto. stimulation. **e,** GABAergic neurons demonstrated closer follow-up activation to USNs compared to non-GABAergic neurons. Y-axis, number of GABAergic vs. non-GABAergic neurons within 3 frames after optogenetic stimulation of USNs. Lag of sampling points indicate the order of frames following USN activation. Lag1 = 100 vs. 0; lag2 = 81.40 vs. 18.60; lag3 = 7.80 vs. 92.2, n=5 animals.

**Extended Data Fig. 13.**
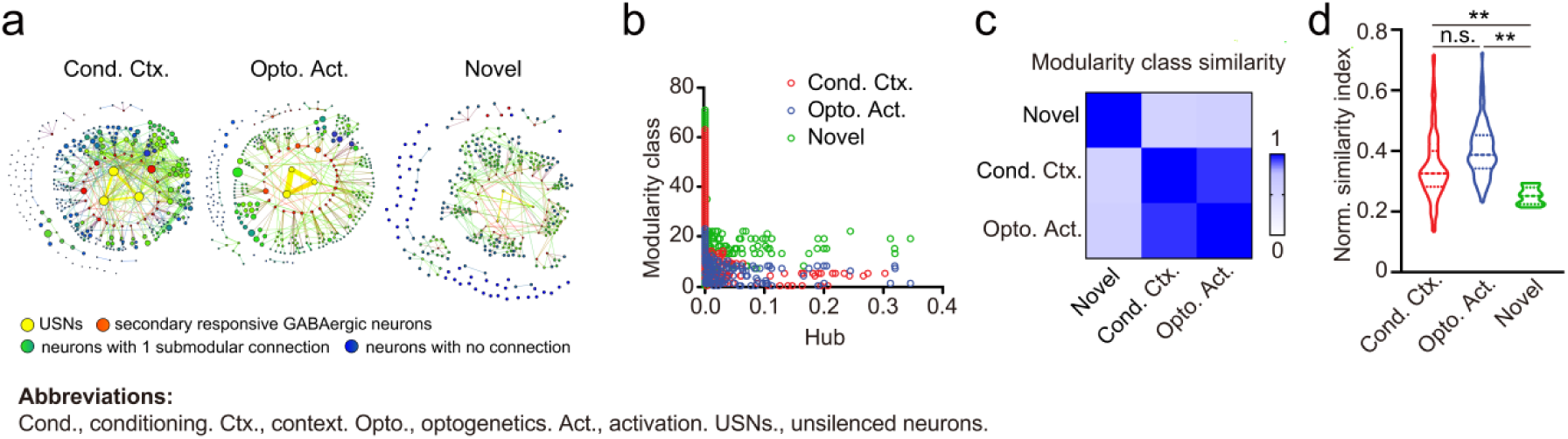
USN activation elicited a disinhibitory hierarchical network relevant for successful retrieval. **a,** Community analysis of neuronal ensembles evoked under various conditions. Optogenetic and contextual stimuli elicit similar multilayer modules, while novel contexts elicit drastically different interneuronal connection strengths and submodular structures. Color gradients and networks illustrate modular patterns between neurons. Node sizes positively correlate to averaged calcium transients of each neuron. **b,** Modularity class distribution of neuronal ensembles activated under different treatments. Modularity classes were calculated from C and denoted community separation. Greater distribution overlap exists between the ensembles elicited by cond. context (red dots) and optogenetic stim. (blue dots) compared to the novel context (green dots). **c,** Modularity class similarity calculation confirmed the highest community similarity under opto. stimulation. and cond. context treatments. The similarity index is calculated by Pearson’s correlation coefficient between distributions of modularity class numbers under respective conditions. n = 330 neurons in 1 animal. **d,** Population distribution of normalized modularity class similarity. Normalization is performed by taking the ratio between each modularity class number and the mean. Cond. Context vs. Opto. stim., *P =* 0.25 *>* 0.05, Cond. Context vs. Novel context: *P =* 0.0004 *<* 0.01, Opto. stim. v.s. Novel context, *P <* 0.01, n = 400 neurons from 4 animals.

**Extended Data Fig. 14.**
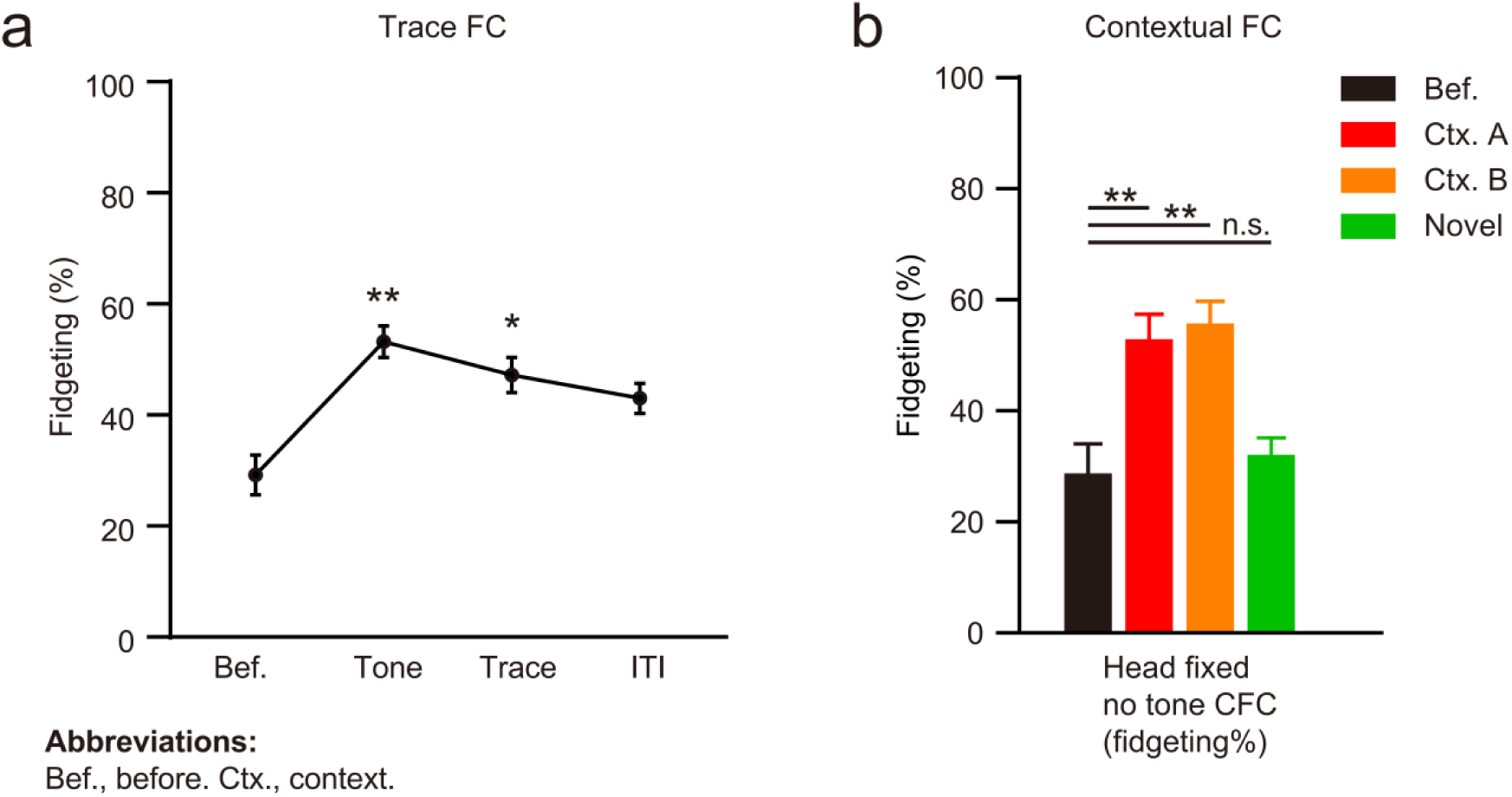
Assessment of paradigm influence on remote memory retrieval. **a,** Successful retrieval of remote fear memory upon tone CS presentation. Results were from the same experimental group reported in Fig. 1D in original submission. Tone vs. before, P = 0.008 < 0.01, Trace vs. before, P = 0.023 < 0.05, ITI vs. before, P = 0.120 > 0.05, n = 5 animals. **b,** Successful retrieval of remote contextual fear memory without tone stimuli under free moving paradigm. Left, freezing response of contextual fear conditioning (CFC) test in free moving conditioning box. Before = 21.00 ± 1.76, Context A = 50.00 ± 2,78, Context B = 52.80 ± 4.44, novel = 23.20 ± 2.25; Context A vs. before, P = < 0.0001, context B vs. before, P < 0.0001, novel vs. before, P = 0.915, n = 5 animals. Right, fidgeting response upon presentation of old context A or old context B. Mice were trained without tone stimuli. Before = 29.00 ± 4.97, Context A = 53.20 ± 4.10, Context B = 56.00 ± 3.61, novel = 32.40 ± 2.60; Context A vs. before, P = 0.0003 < 0.001, context B vs. before, P < 0.0001, novel vs. before, P = 0.935, n = 5 animals.

## Extended Data Tables

**Fig. 1.** Subject information summary.

| Fig. 1 Subject information summary |  |  |  |
| --- | --- | --- | --- |
| Animal ID | Strain | Number of neurons | Number of USNs |
| 1 | C57B6J | 405 | 5 |
| 2 | C57B6J | 258 | 3 |
| 3 | C57B6J | 198 | 2 |
| 4 | C57B6J | 359 | 4 |
| 5 | C57B6J | 502 | 6 |
| 6 | C57B6J | 314 | 4 |
| 7 | C57B6J | 204 | 3 |
| 8 | C57B6J | 193 | 2 |
| 9 | C57B6J | 305 | 3 |
| 10 | C57B6J | 423 | 5 |
| 11 | C57B6J | 333 | 4 |

**Fig. 2.** Subject information summary.

| Fig. 2 Subject information summary |  |  |  |  |
| --- | --- | --- | --- | --- |
| Animal ID | Strain | Number of neurons | Number of GABAergic neurons | Number of USNs |
| 1 | GABA::Ai9 | 220 | 24 | 3 |
| 2 | GABA::Ai9 | 248 | 32 | 3 |
| 3 | GABA::Ai9 | 186 | 17 | 4 |
| 4 | GABA::Ai9 | 302 | 29 | 5 |
| 5 | GABA::Ai9 | 325 | 37 | 5 |
| 6 | GABA::Ai9 | 198 | 35 | 3 |
| 7 | GABA::Ai9 | 586 | 114 | 3 |
| 8 | GABA::Ai9 | 439 | 53 | 4 |
| 9 | GABA::Ai9 | 476 | 69 | 5 |
| 10 | GABA::Ai9 | 625 | 103 | 4 |
| 11 | GABA::Ai9 | 503 | 96 | 5 |

**Fig. 3.** Subjects information summary.

| Fig. 3 Subjects information summary |  |  |  |  |
| --- | --- | --- | --- | --- |
| Animal ID | Strain | Number of neurons | Number of high sync. neurons | Number of USNs |
| 1 | GABA::Ai9 | 220 | 2 | 2 |
| 2 | GABA::Ai9 | 248 | 4 | 3 |
| 3 | GABA::Ai9 | 186 | 4 | 4 |
| 4 | GABA::Ai9 | 302 | 6 | 5 |
| 5 | GABA::Ai9 | 325 | 5 | 5 |
| 6 | GABA::Ai9 | 198 | 3 | 3 |

**Fig. 4.** Two context experiment subjects information summary.

| Fig. 4 Two context experiment subjects information summary |  |  |  |  |
| --- | --- | --- | --- | --- |
| Animal ID | Strain | Number of neurons | Number of USNs for context A | Number of USNs for context B |
| 1 | GABA::Ai9 | 118 | 4 | 3 |
| 2 | GABA::Ai9 | 263 | 3 | 5 |
| 3 | GABA::Ai9 | 194 | 4 | 4 |
| 4 | GABA::Ai9 | 220 | 3 | 3 |
| 5 | GABA::Ai9 | 251 | 4 | 5 |
| 6 | GABA::Ai9 | 252 | 3 | 3 |
| 7 | GABA::Ai9 | 302 | 5 | 2 |

**Fig. 6.** 3D volumetric imaging subjects information summary.

| Fig. 6 3D volumetric imaging subjects information summary |  |  |  |
| --- | --- | --- | --- |
| Animal ID | Strain | Number of neurons | Number of USNs |
| 1 | GABA::Ai9 | 985 | 32 |
| 2 | GABA::Ai9 | 867 | 26 |
| 3 | GABA::Ai9 | 756 | 28 |
| 4 | GABA::Ai9 | 1032 | 31 |
| 5 | GABA::Ai9 | 1135 | 38 |

**Extended Data Fig. 12a.**
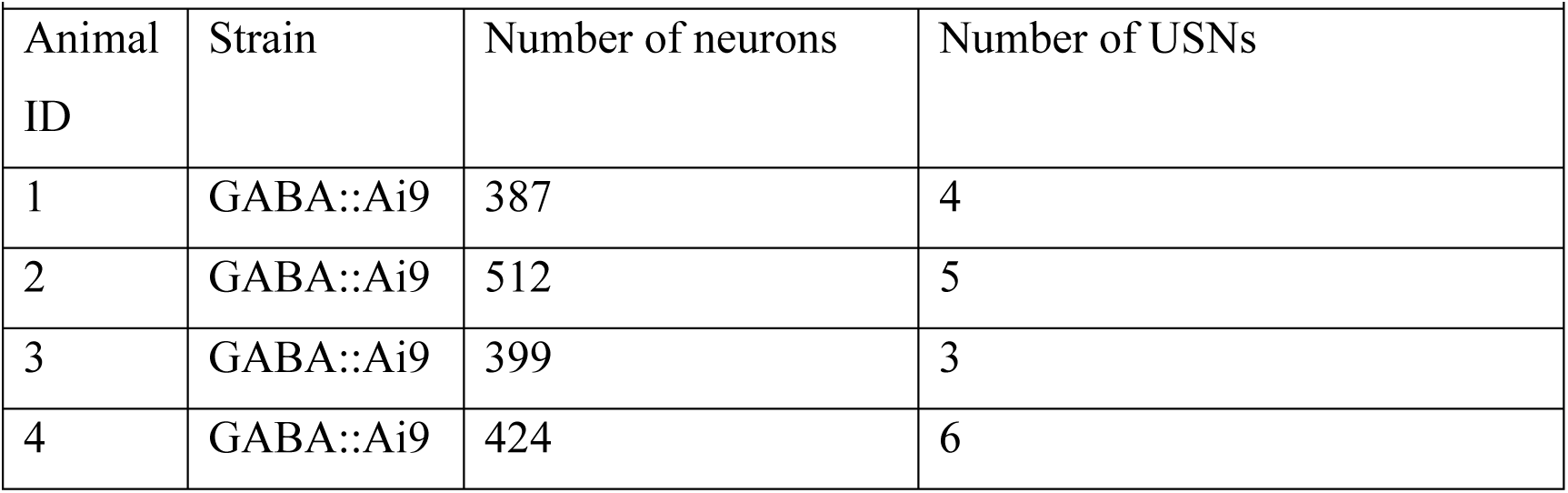
Engram subjects information summary.

